# Peripheral membrane protein endophilin B1 probes, perturbs and permeabilizes lipid bilayers

**DOI:** 10.1101/2024.08.07.606963

**Authors:** Arni Thorlacius, Maksim Rulev, Oscar Sundberg, Anna Sundborger-Lunna

## Abstract

Bin/Amphiphysin/Rvs (BAR) domain containing proteins are cytosolic, peripheral membrane proteins that regulate the curvature of membranes in eukaryotic cells. BAR protein endophilin B1 plays a key role in multiple cellular processes critical for oncogenesis, including autophagy and apoptosis. Amphipathic regions in endophilin B1 drive membrane association and tubulation through membrane scaffolding. Our understanding of exactly how BAR proteins like endophilin B1 promote highly diverse intracellular membrane remodeling events in the cell is severely limited due to lack of high-resolution structural information. Here we present the highest resolution cryo-EM structure of a BAR protein to date and the first structures of a BAR protein bound to nanodiscs. Using neural networks, we can effectively sort particle species of different stoichiometries, revealing the tremendous flexibility of post-membrane binding, pre-polymer BAR dimer organization and membrane deformation. We also show that endophilin B1 efficiently permeabilizes negatively charged liposomes that contain mitochondria-specific lipid cardiolipin and propose a new model for Bax-mediated cell death.

## Main

Regulation of membrane remodeling is essential for maintaining cellular homeostasis.^1^ Proteins from the Bin/Amphiphysin/Rvs-homology (BAR) domain superfamily are peripheral membrane proteins that promote membrane curvature in eukaryotic cells.^2–6^ BAR family members are grouped into classical BAR proteins, N-terminal amphipathic helix-BAR (N-BAR) proteins, BAR-pleckstrin homology (BAR-PH) proteins, Phox homology-BAR (PX-BAR) proteins, Fes/CIP4 homology-BAR (F-BAR) proteins and inverse-BAR (I-BAR) proteins, based on domain organization and function.^3,4^ BAR proteins are typically dimers in solution, where the dimer consists of three α-helices from each monomer that form a 6-helix bundle with a characteristic crescent shape. The concave surface of BAR domains is rich in amino acids with basic side chains, and therefore, preferentially bind anionic membranes via electrostatic interactions.^6–8^ BAR proteins are predominantly associated with helical scaffold assembly and the formation of membrane tubules.^8–26^

Endophilins are a highly conserved group of N-BAR proteins that contain a N-terminal amphipathic helix (H0) and a 20-residue amphipathic insert in helix 1 (H1i).^8,10^ These two motifs contribute to membrane association.^8,10^ A C-terminal Src-homology 3 (SH3) domain, which is connected to the BAR domain by a long flexible linker, mediates protein-protein interactions.^8^ Endophilin B1 preferentially binds to membranes that contain cardiolipin,^25,27^ a mitochondria-specific lipid that plays a critical role in apoptosis.^28–32^ Knockdown of endophilin B1 results in aberrant mitochondrial morphology and a delayed Bax-mediated apoptosis.^28,29^ Evidence suggests that endophilin B1 interacts with Bax via H0.^27,30,31^ Interestingly, the H0 of endophilin B1 is longer than that of other endophilin family members (Table 1) and has a zero net charge, unlike the endophilin A1 H0, which is positively charged.^25,32^ Endophilin B1 has also been shown to interact with Beclin-1 through UVRAG to promote autophagosome formation.^33,34^ During mitophagy, it may interact with mitochondrial inner membrane protein prohibitin and form heterodimers with endophilin B2.^35,36^ Loss of endophilin B1 is seen in several different forms of cancer, which indicates it plays an important tumor suppressor role in the cell.^37–44^

**Table 1.**
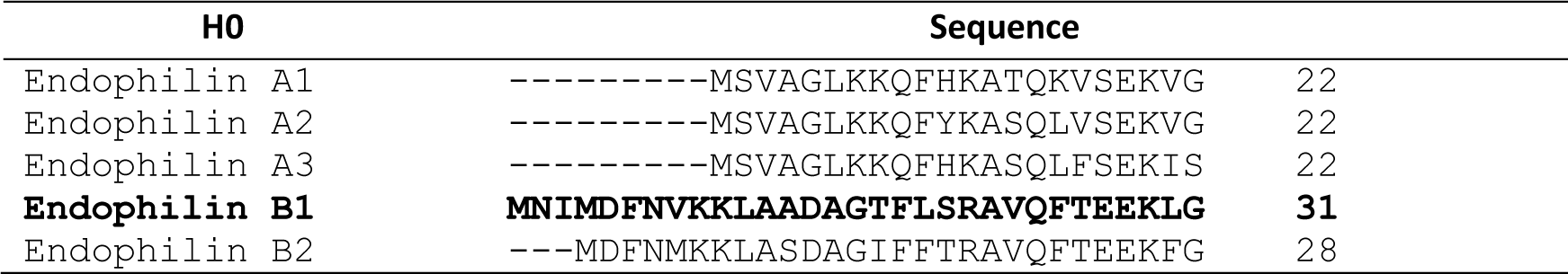
Multiple sequence alignment of endophilin H0 helices.

The exact molecular mechanisms underlying the activity of endophilin B1 at intracellular membranes is unclear. Our previous cryo-electron microscopy (cryo-EM) studies of endophilin B1 reveal it organizes into helical scaffolds on tubulated liposomes that vary greatly in outer diameter (40-60 Å).^25^ This heterogeneity led to poor resolution at the protein-membrane interface, and thus poor insight into the organization of amphipathic regions H0 and H1i.^25^ We further found that the flexible linker-SH3 domain region interacts with H0 in solution and that truncation of the this region yielded more efficient liposome tubulation.^25^ These findings suggest that endophilin B1 is autoinhibited in solution by intramolecular interactions between H0 and the SH3 domain. Similar BAR protein autoinhibition by an SH3 domain was previously proposed for F-BAR protein syndapin-1.^20^

Insight into the structural basis of BAR-mediated membrane remodeling is limited to either static crystal structures of soluble (and often truncated) proteins,^7,10,45–47^ solution NMR structures of small individual domains,^46,48^ or low-resolution cryo-EM maps of helical scaffolds assembled on tubulated liposomes.^16,23–26,49,50^ Furthermore, available crystallographic maps of N-BAR proteins show poor density for amphipathic motifs. This is likely due to the regions assuming helical conformation only when inserted into membranes.^32^ For example, NMR analysis of membrane-bound BIN1 H0 reveals that the N-terminal end is highly flexible, but that the rest adapts a helical conformation.^48^ One available N-BAR crystal structure (BIN2) includes the majority of H0.^51^ Though, the position and orientation of H0 in that model may be called into question as H0 is locked between the BAR domains of symmetry mates in the unit cell. Since the structure contains a large fragment of an H0, it appears to have greatly influenced predictions for other N-BARs, as all AlphaFold models of N-BARs show H0 in a similar position and orientation.^52,53^

Here we present the first single-particle cryo-EM structure of a membrane-bound BAR protein that reaches near-atomic resolution. This allows us to accurately determine the position and variable conformations of amphipathic regions, something that has severely limited previous studies of BAR proteins. Our EM structure of the endophilin B1-nanodisc complex consists of six endophilin B1 dimers bound in two distinct conformations to a single artificial lipid platform. CryoDRGN analysis reveals multiple populations with different stoichiometries, as well as conformational variations of the amphipathic helices and BAR domain of endophilin B1. Structural analyses of full-length endophilin B1 in solution with cryo-EM and SAXS show that the protein is highly flexible. A truncated form of endophilin B1, lacking a SH3 domain has a different shape in solution compared to the full-length protein. These results strengthen our hypothesis that, in solution, the SH3 domain occludes the concave side of the BAR domain, which contributes to autoinhibition. We also present evidence that endophilin B1 permeabilizes liposomes. The phenomenon is dependent on negative surface charge and is enhanced by the presence of cardiolipin. Together, our results present a novel model for BAR protein scaffold assembly and propose a critical role for endophilin B1 in the permeabilization of the outer-mitochondrial membrane during programmed cell death.

## Results

### Endophilin B1 binds to nanodiscs that contain cardiolipin

To understand how endophilin B1 promotes diverse membrane remodeling events in the cell, we wanted to reveal the interaction between endophilin B1 and membranes. Our previous studies show that endophilin B1 organizes into helical polymers on cardiolipin-enriched liposomes in distinct modes.^25^ We proposed that these diverse organizations were regulated by coordinated H0-H1i association with the membrane. However, limited resolution prevented further insight into the exact nature of these interactions. Therefore, to probe the ability of EnB1s amphipathic motifs to drive distinct modes of membrane association and subsequent remodeling, we used the limiting membrane supports of lipid nanodiscs.^54–57^ Nanodiscs have been used extensively to resolve structures of integral membrane proteins,^58^ but only in one occasion to study the structure of a peripheral membrane protein.^59^ We reasoned that these lipid bilayer scaffolds are big enough to allow endophilin B1 binding, but too small to allow polymerization into helical scaffolds.

We generated nanodiscs consisting of membrane scaffolding protein MSP2N2 and lipids with negatively charged headgroups, including cardiolipin. Nanodiscs incubated with endophilin B1 show a substantial shift in molecular weight towards larger assemblies according to size-exclusion chromatography (SEC; Fig. 1a). Incubating nanodiscs with endophilin B1 at different molar ratios (1:2 and 1:10; MSP2N2:Endophilin B1) results in significantly different elution profiles (Extended Data Fig. 1a). Endophilin B1 decoration of nanodiscs was confirmed by negative-stain EM (Extended Data Fig. 1b). Both elution peaks correspond to particles significantly larger than “naked” nanodiscs, and the sample containing more endophilin B1 (1:10) corresponds to a larger particle size than the sample containing less endophilin B1 (Extended Data Fig. 1a). This indicates that higher ratios of endophilin B1 results in nanodiscs with more endophilin B1 decoration.

**Figure 1.**
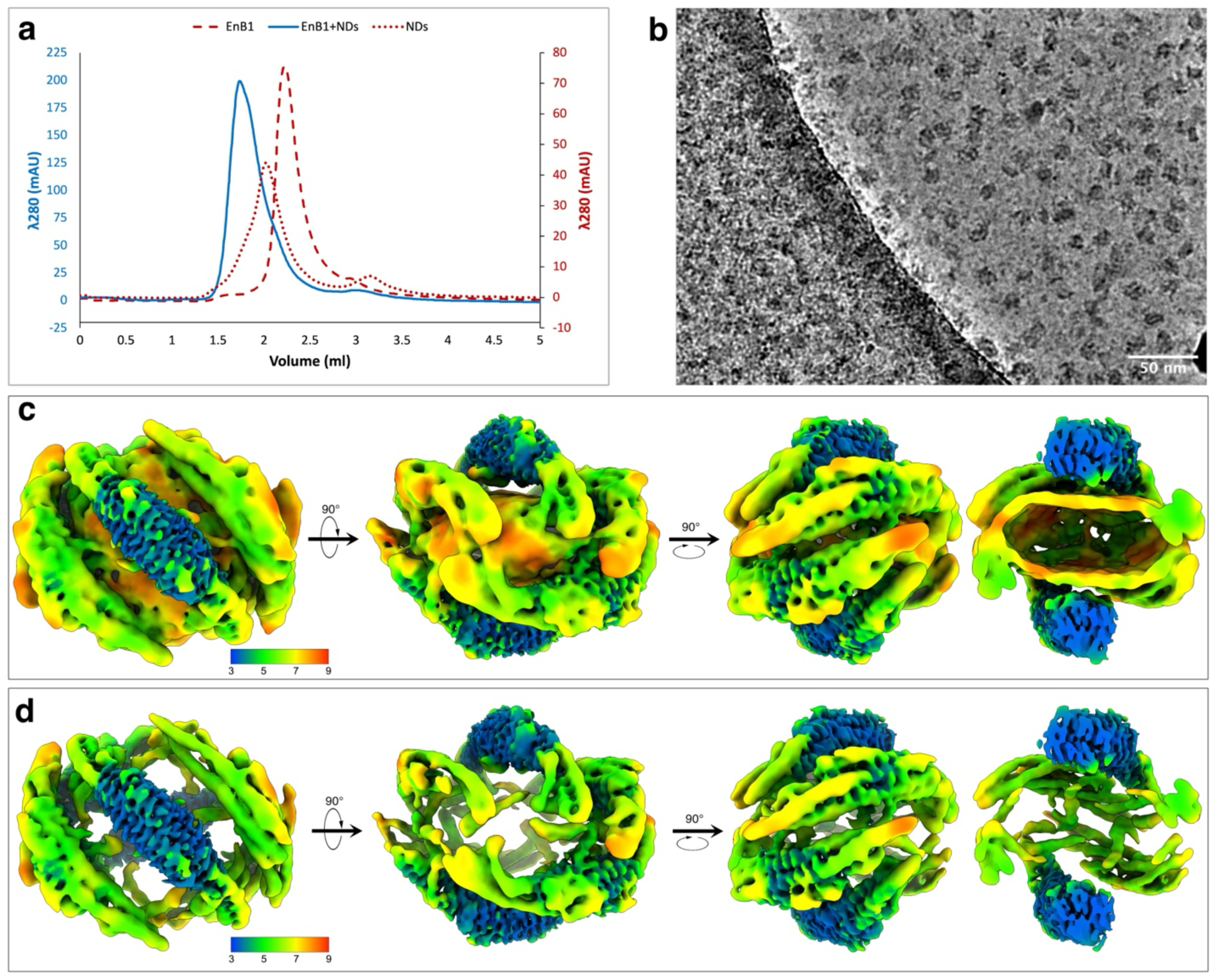
Cryo-EM structure of endophilin B1 (EnB1) bound to a nanodisc (NDs). **a)** Representative SEC runs of EnB1, MSP2N2 NDs and NDs incubated with EnB1 at a molar ratio of 1:10 (MSP2N2:EnB1). **b**) Raw micrograph from high-resolution EnB1-ND data collection. **c-d**) 3D reconstruction of EnB1-decorated NDs at different contour levels showing the local resolution.

### Endophilin B1-decorated nanodiscs with different stoichiometries

Cryo-EM data of the 1:2 molar ratio sample were split into 2 classes. Most particles appeared to consist of empty MSP2N2 nanodisc (Extended Data Fig. 1c) with a smaller subset of nanodiscs decorated with one EnB1 dimer (Extended Data Fig. 1d). Interestingly, the diameter of decorated nanodiscs is roughly 2 nm smaller than that of undecorated nanodiscs. Similarly, nanodiscs decorated with 6 dimers in the 1:10 sample are smaller than undecorated nanodiscs (Extended Data Fig. 1e).

### Amphipathic regions of endophilin B1 anchor to the membrane

The high-resolution data set of endophilin B1-decorated nanodiscs (1:10) contains both conformational and compositional heterogeneity, despite eluting as a single SEC peak (Fig. 1a and 1b). Electron density maps consisting of >5 membrane-bound endophilin B1 dimers were improved iteratively during several rounds of multi-class heterogeneous *ab-initio* reconstruction followed by heterogeneous refinement in cryoSPARC (for the complete workflow see Extended Data Fig. 2).^60^ The final volume consisted of 6 endophilin B1 dimers and had a nominal resolution of 3.88 Å (Extended Data Fig. 3a, Extended Data Table 1). Strong density could be observed for 4 dimers and slightly weaker density for two additional dimers. The volume contains a membrane bilayer density with anchored amphipathic helices (Fig. 1b). Interestingly, at higher contour levels the density corresponding to the membrane bilayer disappears, and where we expect to observe density corresponding to MSP2N2 scaffolds, there is none (Fig. 1c). Instead, we observe density that belongs to endophilin B1 amphipathic helices.

The local resolution of the BAR domains is higher than that of the inserted amphipathic helices, suggesting the protein is flexible at the protein-lipid interface (Fig. 1c). The highest local resolution is found at the center of the BAR domains, at the dimerization interface, however the distal ends of the BAR domains appear flexible.

### Endophilin B1 amphipathic regions undergo major conformational changes upon membrane binding

Together, six dimers form a cage (“mini scaffold”) around a patch of bilayer, where each dimer makes direct contact with neighboring dimers through their respective H0 and H1i motifs (Fig. 1c). Dimers bound to the surface of the bilayer and dimers that straddle the sides of the nanodisc have distinct amphipathic helix conformations (Fig. 2a and 2b). These different classes of dimers will be referred to as “center” and “side” dimers, respectively.

**Figure 2.**
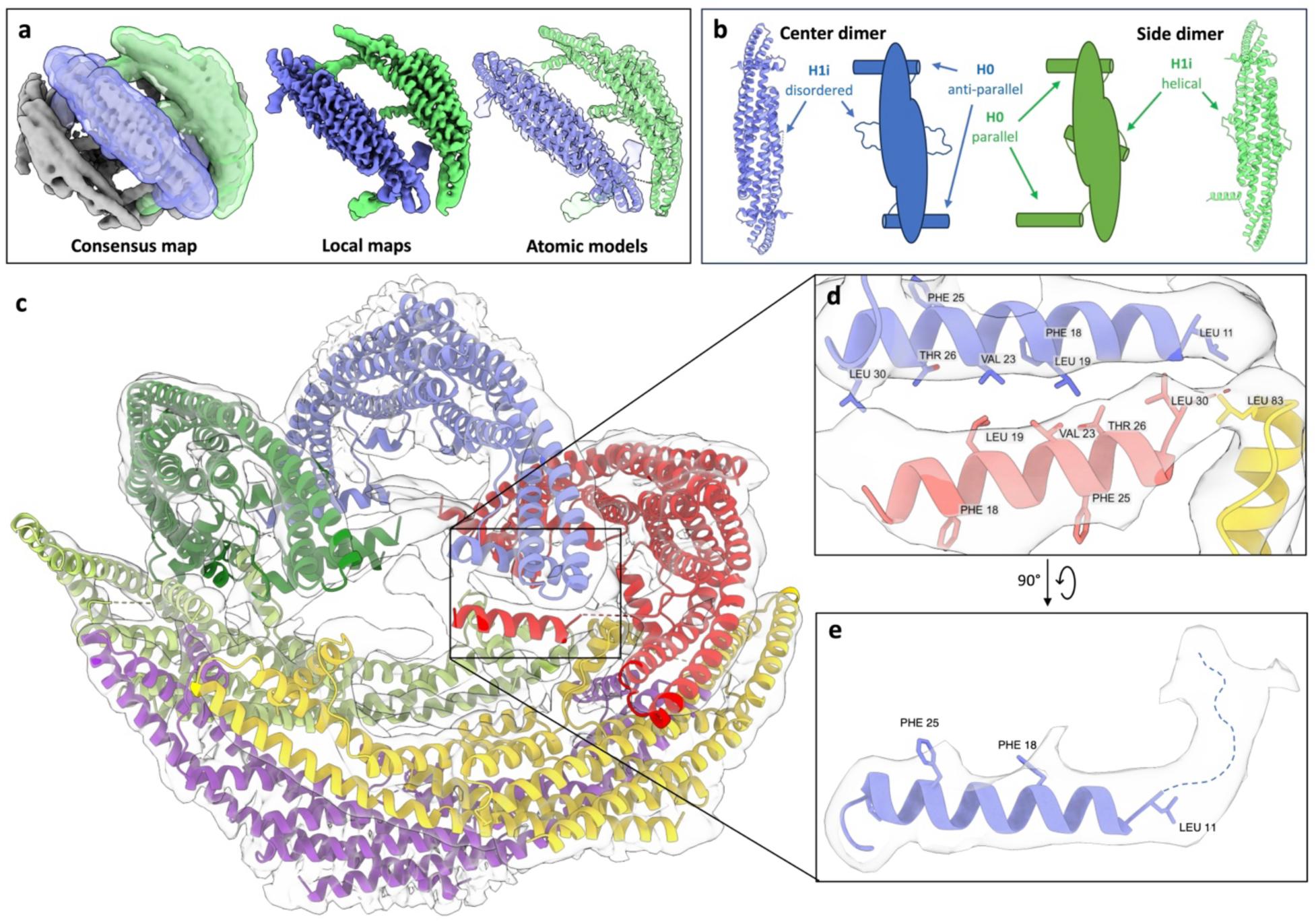
Focused refinement and atomic models of membrane-bound EnB1. **a**) Left: Electron density map of EnB1 on a ND with masks for one center dimer and one side dimer. Middle: Resulting local refinement maps of the center and side dimers. Right: Atomic models docked into both density maps. **b**) Top views of the atomic models and cartoon representations highlighting the different conformations of the amphipathic helices. **c**) Models for all dimers docked into the original density map. **d**) Interactions between multiple amphipathic motifs. **e**) The N-terminus of H0, is flexible in the structure. The dotted line represents the flexible N-terminus in the unmodeled part of the H0 electron density.

In center dimers, H1i is predominantly disordered. At lower contour levels the disordered region appears close to the BAR domain, brushing the membrane surface, but not inserted. H0s are oriented anti-parallel in relation to each other. In side dimers, both helical and loop regions of H1i have visible density and H1i is inserted into the membrane. Interestingly, H0 are oriented parallel to each other. Locally refining the classes individually resulted in reconstructions of 3.45 Å (center dimer; Extended Data Fig. 3b) and 3.60 Å (side dimer; Extended Data Fig. 3c), respectively. These two maps were used to build atomic models (Fig. 2b, Extended Data Table 1).

The atomic models consist of residues Leu11-Leu252. Residues Met1-Lys10 of H0 were omitted as weak density indicates that the N-terminus is disordered (Fig. 2e), while residues L11-L30 assume a helical conformation. As H1i is disordered in the center dimer class, residues Glu76-Ile96 were deleted from the model (Extended Data Fig. 3d). However, there is clear density corresponding to the whole H1i including the loop connecting H0 to the BAR domain in the side dimer reconstruction (Extended Data Fig. 3e).

The H2-H3 hinge region is flexible when membrane-bound The BAR domain H2 can be split into two segments. The portion that contributes to dimer formation is more rigid, which is reflected in higher local resolution in the EM maps for that section (Extended Data Fig. 3f and 3g). The portion that extends away from the dimer interface has significantly worse density, indicating that there is conformational heterogeneity present. The residues between these segments form a hinge that moves in concert with the hinge region that is also present in H3. The key residues that appear to facilitate movement are two glycines, Gly153 and Gly215, that nick their respective helices, splitting them into rigid and more flexible sections (Extended Data Fig. 3h). Docking the models into the original EM map (Fig. 2c) suggests that inter-dimer contact sites consist of hydrophobic interactions between leucines and valines (Leu19, 30, 83 and Val23) (Fig. 2d).

### Neural network analysis reveals side-to-side assembly and distortion of nanodisc shape

The workflow described above yielded two 3D reconstructions of near-atomic resolution that were used to build atomic models. However, we were also interested in assemblies with different stoichiometries that had been filtered away during the refinement process. To untangle the compositional and conformational heterogeneity present in the sample, we used cryoDRGN.^61–63^ Two groups of particles, one, which resulted in a high-resolution reconstruction, and another, further upstream in the cryoSPARC workflow (maps * and †, respectively, in Extended Data Fig. 2), were exported to cryoDRGN. Training results for the high-resolution reconstruction reveal that a shift in H0 density is coupled with conformational changes in the side dimer BAR domain (Movie 1). Principle component (PC) analysis (Fig. 3a) shows that the shape of the nanodisc changes and becomes more elliptical as the distance between the distal tips of side dimers on opposite sides of the platform increases by ∼1-2 nm (Movie 2, Fig. 3). As the nanodisc becomes more elliptical, the center dimers H0s move closer together and the membrane density becomes more distinct. CryoDRGN heterogeneous *ab-initio* reconstructions of the larger group of particles effectively sorted particles with different stoichiometries (Fig. 4a, Movie 3). 3D reconstructions were generated for nanodiscs classes with 3-6 endophilin B1 dimers, at nominal resolutions between 6.40 Å and 9.59 Å (Fig. 4b-k). These reveal how endophilin B1 mini scaffolds assemble side-to-side on nanodiscs.

**Figure 3.**
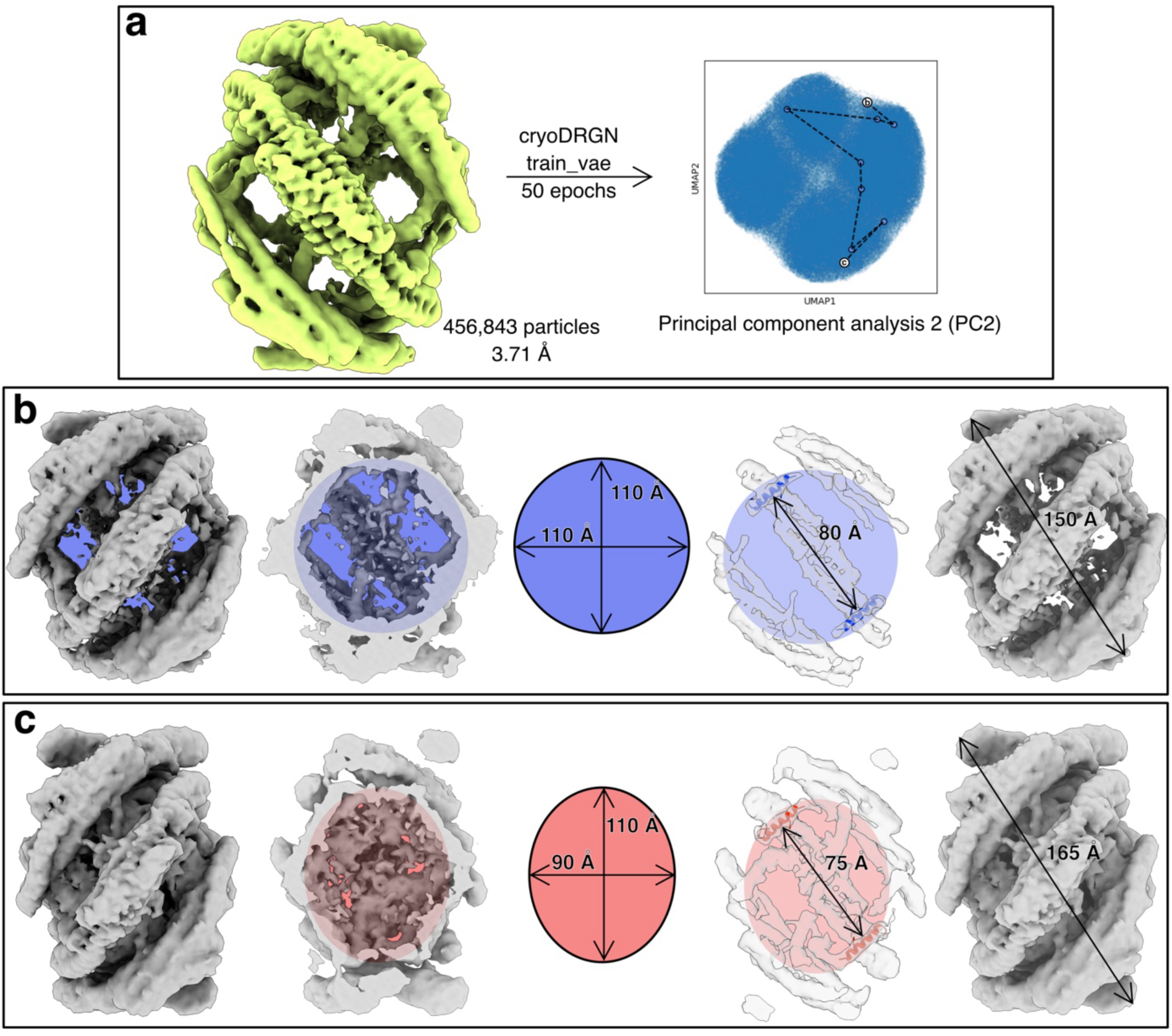
CryoDRGN predicts that EnB1 distorts the shape and affects the membrane fluidity of NDs. **a**) 3D reconstruction of particles used for analysis with cryoDRGN train_vae and the resulting UMAP distribution showing traversal along principal component 2 (PC2 traversal can be seen in Movie 3). Volumes generated by cryoDRGN representing the minima (**b**) and maxima (**c**) of PC2. CryoDRGN predicts a more intense signal from the lipid bilayer in (**c**) compared to (**b**).

**Figure 4.**
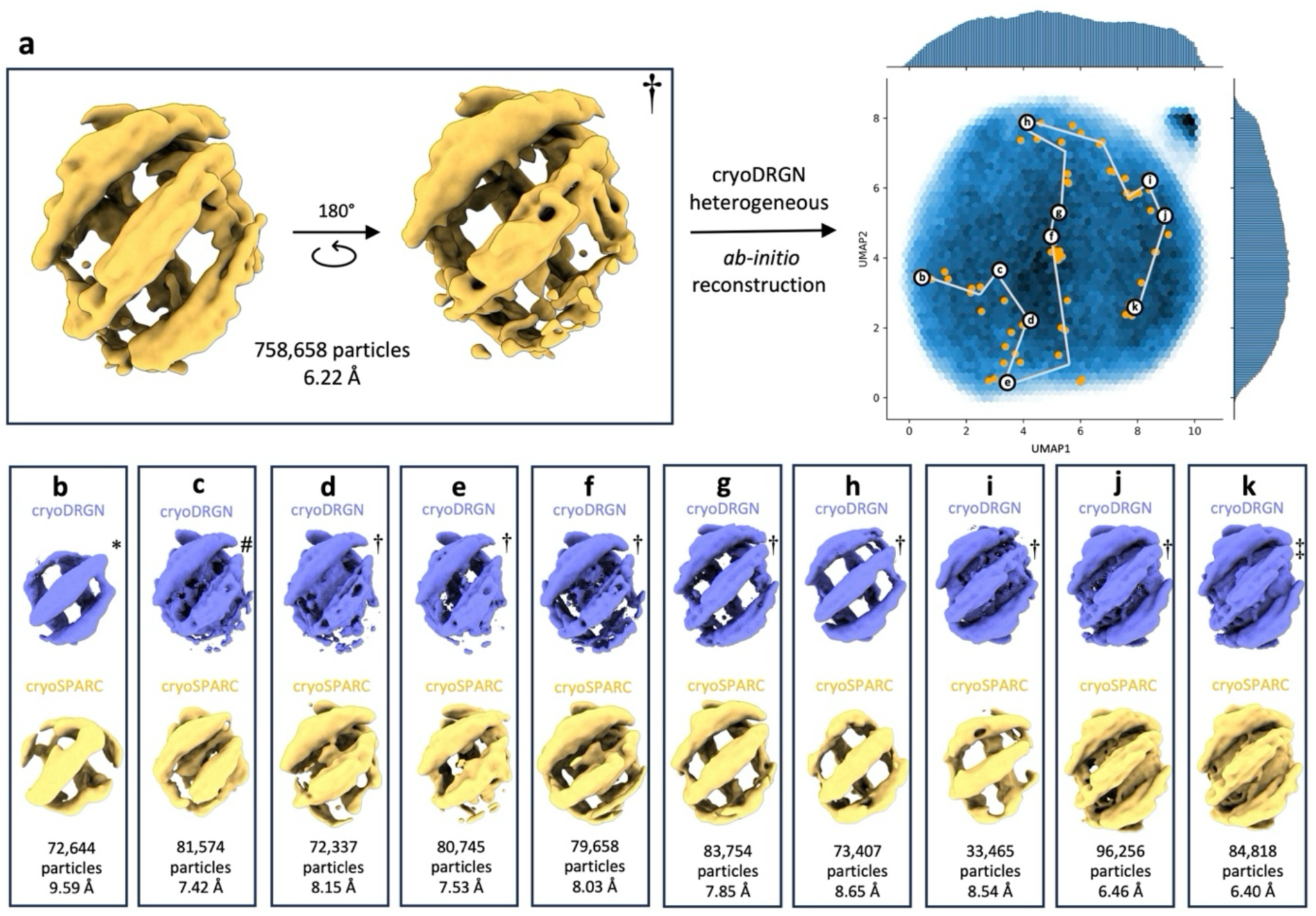
CryoDRGN reveals side-to-side assembly of EnB1 on NDs. **a**) 3D reconstruction of particles used in cryoDRGN heterogeneous *ab-initio* reconstruction and the resulting UMAP distribution showing locations of different k-means clustered classes (**b**-**k**). Symbols mark the different stoichiometries present in each class of particles: (*) three, (#) four, (†) five, or (‡) six dimers bound per ND.

### The linker-SH3 domain region is flexible when endophilin B1 is membrane-bound

We have previously shown that the SH3 domain negatively regulates the ability of endophilin B1 to mediate membrane tubulation, suggesting it plays an important role in the regulation of endophilin B1 membrane activity.^25^ In solution, the SH3 domain associates with H0. This observation led us to propose that, in solution, the SH3 domain is fixed in position near the concave surface of the BAR domain by H0, and that H0 association with membranes regulates its conformation, and subsequently, the ability of endophilin B1 to cause membrane curvature. Previous studies of endophilin A1 propose that the SH3 domain may bind opposite ends of the BAR dimer.^23,64^ Interestingly, no density corresponding to the SH3 domain-linker region is visible in any of our maps, either near the concave surface or proximal to the lateral part of the BAR domain. This indicates that this region is highly flexible when endophilin B1 is membrane-bound. To determine whether membrane binding regulates the organization of this flexible region, we attempted to determine the structure of the cytosolic, i.e., the soluble state of endophilin B1 using cryo-EM. However, vitrified grids consistently had poor ice quality and particle distribution, with most adhering to the carbon, to the edge of holes or clustering in discrete patches (Extended Data Fig. 4a and 4b). Low concentration of protein (5 μM) yielded slightly better ice, which allowed collection of a small data set and generation of 2D class averages (Extended Data Fig. 4c). Ultimately, no useful 3D reconstruction could be obtained due to preferred particle orientation.

Small-angle X-ray scattering (SAXS) data was collected using soluble endophilin B1 at three different concentrations (4 μM, 8 μM and 16 μM; Extended Data Fig. 5a-c and 6a). We observe differences in the radius of gyration (*R_G_*) and the polydispersity of the protein at different concentrations (Extended Data Fig. 6b). At higher concentrations (8-16 μM), endophilin B1 is primarily dimeric and partly disordered, whereas at low concentration (4 μM), it is monomeric and mostly disordered (Extended Data Fig. 6c-e). Analysis of SAXS data of truncated endophilin B1 lacking the SH3 domain (endophilin B1_ΔSH3; Extended Data Fig. 5d and 7b) indicates that the mutated protein is more globular than the wild type form (Extended Data Fig. 7c-f). We observe a decrease in *R_G_* compared to the wild type. The SAXS *ab-initio* reconstructions of wild type endophilin B1 and endophilin B1_ΔSH3 show BAR domains with distinct shapes. The reconstructions are similar in length; however, the wild type reconstruction has added density on the concave side of the BAR domain (Extended Data Fig. 7f). Endophilin B1_ΔSH3 and full-length endophilin B1 have similar elution volumes in SEC despite a predicted ∼12 kDa difference in MW (Extended Data Fig. 7a).

### Endophilin B1 permeabilizes membrane vesicles

It has been previously shown that insertion of amphipathic regions can cause membrane permeabilization.^65–69^ We observe variability in the membrane bilayer upon insertion of H0 and H1i. To determine whether endophilin B1-mediated membrane disruption drives membrane permeabilization, we added endophilin B1 to liposome with encapsulated quenched calcein (Table 2). We find that addition of endophilin B1 to these membrane vesicles lead to significant increase in calcein fluorescence as a result of membrane permeabilization and de-quenching (Fig. 4). Interestingly, the ability of endophilin B1 to promote membrane permeabilization is dependent on the lipid composition. Endophilin B1 shows low permeabilization activity when added to liposomes with a net neutral charge (Mix 1) and significantly higher activity when added to net negatively charged liposomes (Mix 2) (Fig. 5a). We observe significantly more permeabilization when vesicles were prepared with the addition of phospholipids that contribute to membrane packing defects (Mix 3).^70,71^ This effect is further increased when cholesterol is removed (Mix 4). Robust permeabilization of vesicles was observed when a higher concentration of endophilin B1 (500 nM vs 5 μM) was added to vesicles with 20% CL (Mix 5) (Fig. 5b). The most efficient permeabilization is observed at 37 °C.

**Figure 5.**
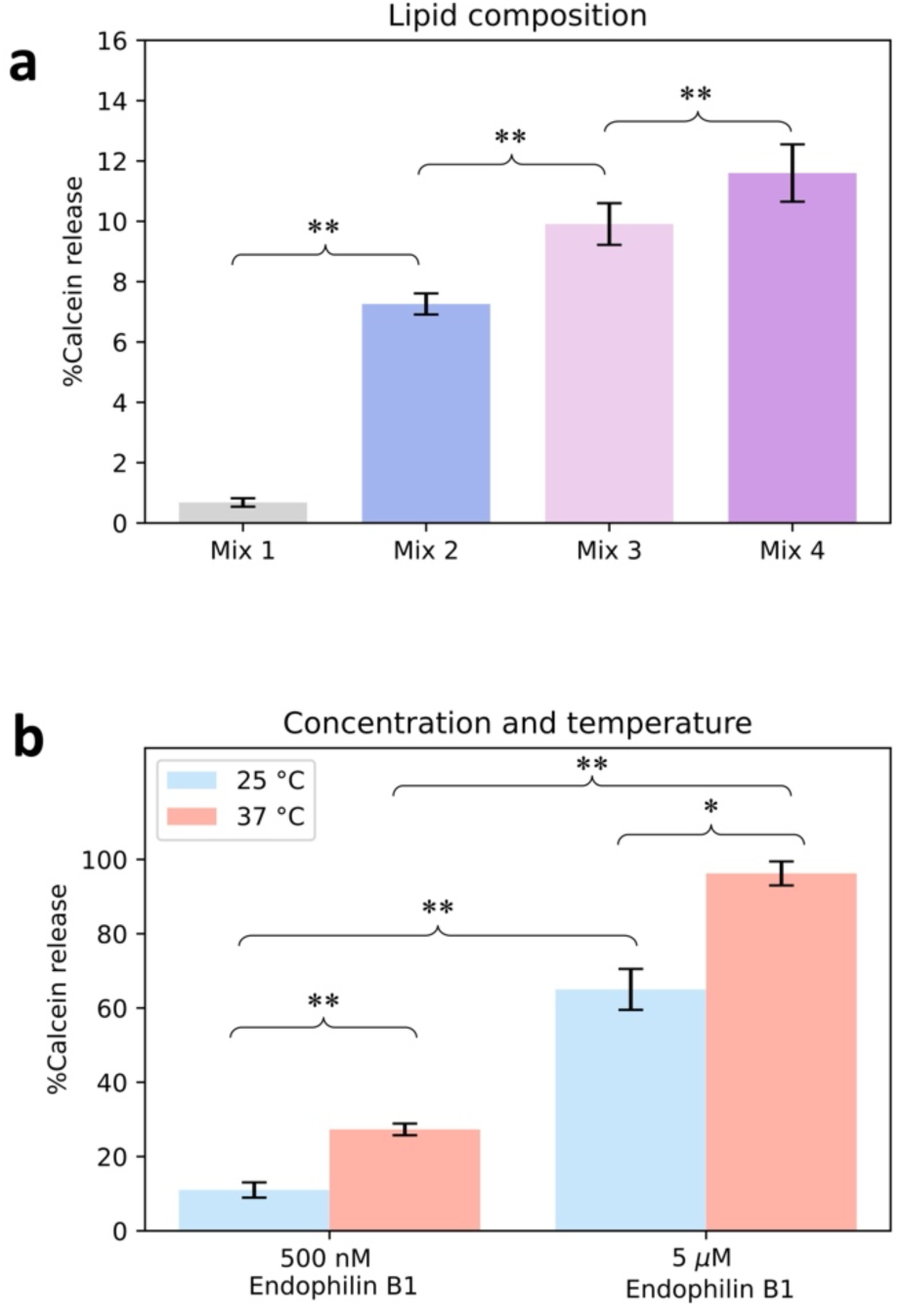
EnB1-mediated liposome permeabilization is dependent on membrane lipid composition. Bar graphs showing % calcein release from liposomes with different lipid compositions (Table 2) after addition of EnB1 (**a**) and % calcein release from liposomes containing lipid mix 5 after addition of increasing concentrations of EnB1 at RT and physiological temperature (**b**). Data represents mean ± standard deviation, ** p < 0.05, *** p < 0.01.

**Table 2.** Permeabilization assay using endophilin B1 and liposomes of different lipid compositions (lipid ratio shown as molar percentage) where *n* is the number of technical replicates. Results are shown as the mean ± standard deviation of the mean.

| Name | Lipid composition | Protein conc. (μM) | Temperature ($^{\circ}$C) | $n$ | Calcein release (%) |
| --- | --- | --- | --- | --- | --- |
| Mix 1 | 100% DOPC | 0.5 | 25 | 18 | 0.68 ± 0.14 |
| Mix 2 | 55% DOPS, 40% DOPE, 5% Cholesterol | 0.5 | 25 | 18 | 7.3 ± 0.4 |
| Mix 3 | 50% DOPS, 40% DOPE, 5% Cholesterol, 5% 18:1 CL | 0.5 | 25 | 18 | 9.9 ± 0.7 |
| Mix 4 | 55% DOPS, 40% DOPE, 5% 18:1 CL | 0.5 | 25 | 18 | 12 ± 1 |
| Mix 5 | 40% DOPS, 40% DOPE, 20% 18:1 CL | 0.5 | 25 | 18 | 11 ± 2 |
| | | 0.5 | 37 | 9 | 27 ± 2 |
| | | 5 | 25 | 6 | 65 ± 6 |
| | | 5 | 37 | 6 | 96 ± 3 |

## Discussions

Endophilin B1 binds to nanodiscs with a stoichiometry proportional to the molar ratio between endophilin B1 and MSP2N2. The 1:10 molar ratio cryo-EM data set contained substantial compositional and conformational heterogeneity. Through saturation of the available lipid surface area during sample preparation and extensive cryo-EM data processing of a large data set, a near-atomic resolution reconstruction could be produced for particles with 6 endophilin B1 dimers bound per nanodisc. These structures are the highest resolution EM structures of a BAR protein to date. Another benefit to this approach for studying peripheral membrane proteins is that it can capture the shape and orientation of membrane-bound amphipathic regions.

### Endophilin B1 amphipathic regions displace MSP scaffolds

Initially, we suspected that the amphipathic helices of endophilin B1 might cling to MSP2N2. Surprisingly, we found no density for MSP2N2 in our maps, despite its presence in the sample confirmed by Western Blot analysis (Extended Data Fig. 9). Only amphipathic regions of endophilin B1 are present in our final map. We speculate that destabilization of the nanodisc lipid bilayer by endophilin B1 causes the displacement of MSP2N2. It is replaced by the amphipathic regions of endophilin B1, which stabilize the nanodisc bilayer. This would mean that our MSP nanodiscs are truly endophilin B1-decorated bicelles. There are multiple reports of proteins and peptides (in addition to MSPs) that can form discoidal lipoprotein particles.^72–75^ Perhaps isolated endophilin H0s could represent another method of creating nanodiscs-like particles similar to the Salipro system.^73^

### Endophilin B1 amphipathic regions drive heterogeneous assembly

The endophilin B1 dimers decorating the discs could be split into two categories based on where they bound to the nanodiscs; either on the flatter side of the disc (center) or on the more curved edge of the disc (side). Focused refinement revealed that the major differences between these two dimers are the conformations of their respective amphipathic motifs. H0 is membrane-bound in both dimer categories, but H1i is not, which indicates that H0 is responsible for initial membrane binding and that H1i insertion occurs later. In the consensus structure, we observe that intermolecular interactions occur on the membrane surface through amphipathic motifs (Fig. 2c). These motifs all bind to or near the nanodisc edge, where local curvature is the highest. H0s organize anti-parallel to each other. The loop that links H0 to the BAR domain is flexible enough to accommodate H0 twisting ∼180°, allowing side-to-side assembly. Diverse orientations of H0 positioning could explain why endophilin B1 (and other endophilins) produce heterogeneous helical scaffolds,^23,25,49^ as there are multiple plausible ways it could oligomerize to form scaffolds (Fig. 6a).

**Figure 6.**
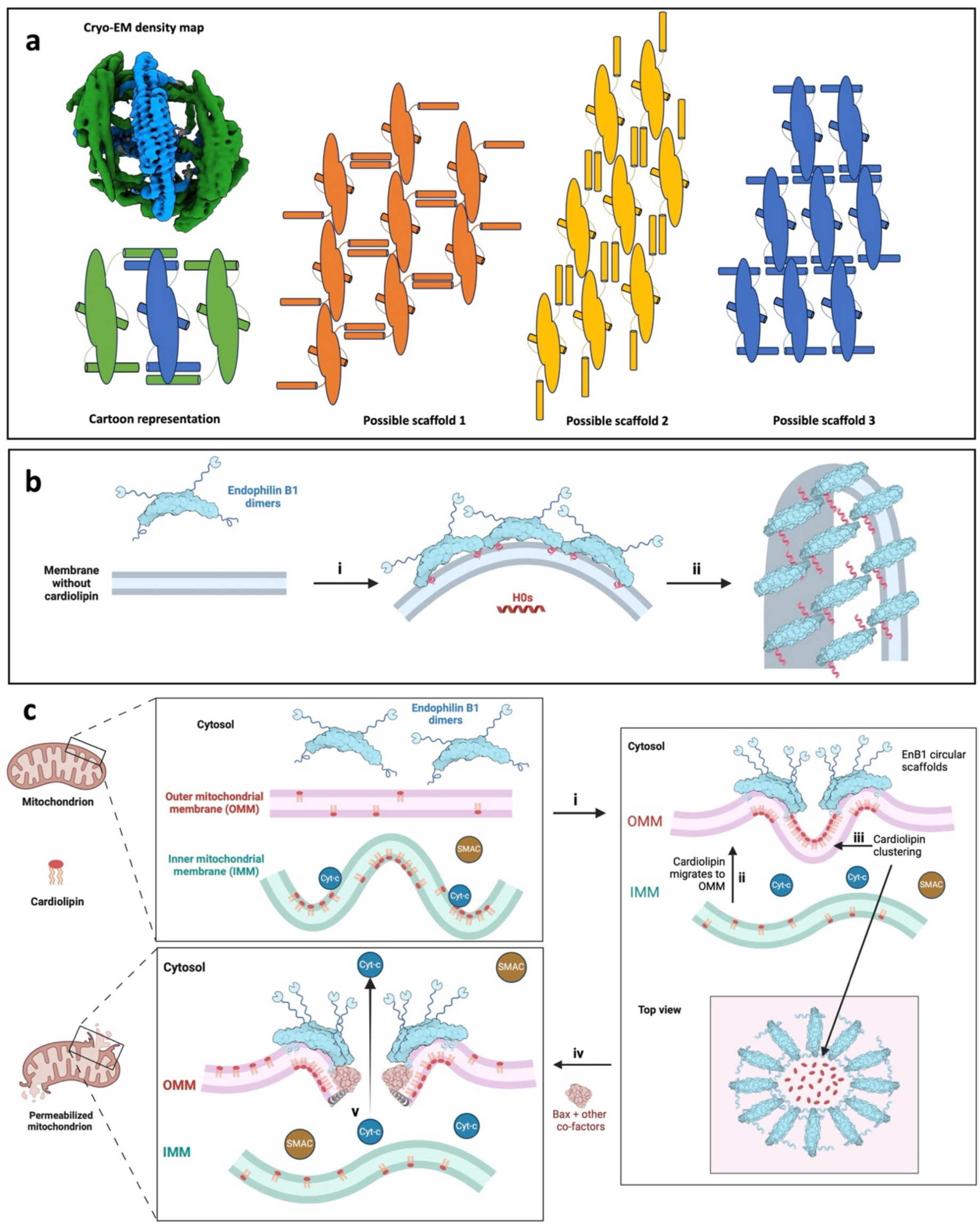
Amphipathic motif assembly promotes EnB1 activity at outer-mitochondrial membranes (OMMs) and drives apoptosis. **a**) Center and side dimers colored in the consensus map and a cartoon representation. Three potential forms of helical scaffolds with different orientations of H0. **b**) Upon binding to a membrane that does not contain cardiolipin, EnB1 oligomerizes end-to-end (i) to form helical scaffolds that enforce membrane tubulation (ii). **c**) Following apoptotic stimuli (i), cardiolipin (a non-bilayer lipid that induces negative curvature) migrates to the OMM (ii). The inner mitochondrial membrane (IMM) is destabilized leading to cytochrome-c (Cyt-c) dissociation. EnB1 binds the OMM and oligomerizes side-to-side into circular scaffolds that cause local negative curvature and further clustering of cardiolipin (iii). Bax and other co-factors bind cardiolipin-rich sites (iv). Bax oligomerization induces leakage of Cyt-c from the intermembrane space (v), which triggers a proteolytic cascade, culminating in cell death.

### BAR domain flexibility at the membrane surface

There was unexpected variability in the conformations of helices 2 and 3 at the tips of the BAR domain. CryoDRGN results show that the BAR domain is flexible and can assume and induce different levels of curvature. This is also apparent when the two atomic models are compared. This is mainly due to kinks in helices 2 and 3 that accommodate the hinge movement. Kinks, and therefore possible hinge movement, are present in other BAR proteins, such as sorting nexins, FCHo and syndapin.^4^ In the H2 and H3 kinks of endophilin B1 there are two glycines, Gly153 and Gly215. These residues are conserved in endophilins B1/2, but not in the endophilin A subfamily, suggesting that endophilin B1 and B2 can adapt to and induce a wider range of curvature than endophilins in the A subfamily. This further supports our notion that endophilin B1 is uniquely capable of promoting diverse membrane remodeling events.

### Intramolecular binding involving the linker-SH3 region

Our previous data suggests that the SH3 domain autoinhibits the membrane remodeling activity of endophilin B1.^25^ Interestingly, we observe no SH3 domain density in our maps of membrane-associated endophilin B1 (Fig. 1b and 1c). Furthermore, attempts to determine the binding location of the SH3 domain in the soluble form of the protein were inconclusive (Extended Data Fig. 4). Soluble endophilin B1 is highly flexible (Extended Data Fig. 6c), indicating that the linker-SH3 domain region adopts more than one conformation in solution. However, wild type endophilin B1 does contain additional density on the concave side of the protein, between the H0s (Extended Data Fig. 6e). We speculate that this density constitutes the SH3-linker region, and that the intramolecular interactions are dynamic. Interestingly, removal of the SH3 domain renders endophilin B1 less flexible (Extended Data Fig. 7d). This indicates that these intramolecular interactions are more prevalent without the SH3 domain. We propose that the linker is responsible for mediating intramolecular interactions, and that these interactions are weakened by the presence of the SH3 domain.

### Monomeric endophilin B1 is polydisperse

Endophilin B1 is monomeric in solution, as evidenced by 2D classes, where only three BAR domain helices can be observed (Extended Data Fig. 4), and SAXS analysis (Extended Data Fig. 6b). Furthermore, monomeric endophilin B1 is more polydisperse than its dimeric form. This is corroborated by observations that dimerization decreases the flexibility of endophilin B1. This may be due to the stabilization of the alpha helices that constitute the dimer interface upon assembly. It is also possible that intramolecular interactions involving the linker-SH3 domain are weaker in the monomeric form, which would increase polydispersity.

### Endophilin B1 perturbs and permeabilizes membranes

We observe stronger density for the lipid bilayer when the BAR domain “flexes”, i.e. brings the H0s of individual endophilin dimers closer together (Movie 2). This indicates that endophilin B1 mini scaffold assembly greatly disturbs the membrane bilayer. Interestingly, we also observe assembly of endophilin B1 mini scaffold on large cardiolipin-containing membrane liposome (data not shown), suggesting that this form of endophilin B1 assembly occurs even when space is not limiting. Both endophilin scaffolds and isolated H0s limit lipid diffusion, which has been suggested to cause vesiculation.^76,77^ It was previously reported that endophilin B1 induces minor leakage of fluorescein isothiocyanate-labeled dextran loaded liposomes.^27^ In this study, we show that endophilin B1 effectively permeabilizes mitochondria-like liposomes and that this activity is dependent on the membrane lipid composition, specifically the presence of cardiolipin, temperature and protein concentration (Fig. 4). Moreover, we present the structural basis for endophilin B1-mediated membrane remodeling, revealing how H0 and H1i insertion and interactions cause destabilization.

### The role of Endophilin B1 during intrinsic apoptosis

Endophilin B1 notably has the longest H0 of all endophilins. In our electron density maps, the H0 residues suggested to associate with Bax do not assume a helical conformation (as predicted by AlphaFold, entry: Q9Y371). Instead, H0 is helical close to its C-terminus, but unwound towards the middle while the N-terminus is flexible (Fig. 2e). Similar flexibility of membrane-bound H0 has also been observed in BIN1.^48^ We find that the N-terminus probes the membrane surface. We suspect that this flexibility is critical for the role of endophilin B1 at OMMs, and that H0 not only targets endophilin B1 to the OMM, but also uniquely interacts with Bax during OMM permeabilization and apoptosis.

Helical scaffold formation by BAR proteins is well established.^3,4,6^ A well-studied example is that of endophilin A1 during endocytosis. When endophilin A1 binds to PIP_2_-rich sites on the inner leaflet of the plasma membrane and oligomerizes, it stabilizes positive curvature by the formation of a helical scaffold (Fig. 5a).^78,79^ The SH3 domain then recruits effector proteins, including dynamin 1, which promotes membrane scission and helical scaffold disassembly.^22^ Endophilin B1 shows no preference for PIP2 but rather cardiolipin and PA (data not shown). However, it has not been determined how endophilin interacts with membranes rich in cardiolipin, a cone-shaped lipid, which favors negative curvature, such as the outer mitochondrial membrane (OMM).^70,80,81^

Following apoptotic stimuli and increased generation of reactive oxygen species, oxidized cardiolipin migrates to the OMM (Fig. 5b), ^82–84^ where it may act as a recruitment platform for Bax.^85^ We propose that during apoptosis, endophilin B1 localizes to cardiolipin-rich areas on the OMM, where is stabilizes the negative curvature generated by cardiolipin clustering. As a result of the local membrane curvature, endophilin B1 assembly is limited to circular mini scaffolds, where dimers interact side-to-side instead of end-to-end. Further clustering of cardiolipin into pits is stabilized by the mini scaffolds limiting lipid diffusion. This creates areas of highly unstable membrane patches onto where Bax and other pro-apoptotic factors are recruited. Once they insert into the membrane and oligomerize, OMM permeabilization, cytochrome-c leakage and ultimately cell death, is inevitable.

In summary, we show multiple features of the structural basis for endophilin B1 membrane disruption, which on a membrane rich in packing defects and negative charge, leads to permeabilization. We propose that endophilin B1 and cardiolipin coordinate to promote Bax-mediated cell death through an interplay of scaffold assembly, negative curvature and chaotic insertion of amphipathic regions.

## Acknowledgements

Data was collected at the Cryo-EM Swedish National Facility funded by the Knut and Alice Wallenberg, Family Erling Persson and Kempe Foundations, SciLifeLab, at Cryo-EM Uppsala, and at the Karolinska Institutet 3D-EM facility. 3D reconstruction was performed on the DaVinci cluster with generous technical assistance from Professor Filipe Maia and Dr Daniel Larsson. CryoDRGN computation was enabled by the Berzelius resource provided by the Knut and Alice Wallenberg Foundation at the National Supercomputer Centre. We also acknowledge the European Synchrotron Radiation Facility (ESRF) for provision of synchrotron radiation facilities and we would like to thank Dr. Petra Pernot for assistance and support in using beamline BM29. We would like to thank the Swedish Research Council (2021-05-423 ASL) for funding this project.

## Methods

### Expression and purification of endophilin-B1

His_12_-SUMO-endophilin-B1, both full-length and truncated forms, were expressed recombinantly in *E. coli* and purified as described previously.^25^ Size-exclusion chromatography (Extended Data Fig. 8a) followed by SDS-PAGE analysis was used to estimate the purity of the sample (Extended Data Fig. 8b). Pure endophilin B1 (≥99% w/w) was concentrated to 2 mg/ml, flash frozen in liquid N_2_ and stored at -70 °C until use.

### Expression and purification of MSP2N2

His_6_-MSP2N2 (Addgene plasmid #29520)^86^ was transformed into BL21(DE3) cells and expressed and purified as previously described with some modifications.^87^ Bacteria were grown in terrific broth (TB) and induced with 0.2 mM IPTG at OD_600_ = 0.5, followed by expression at 37 °C for 3 hours. Cells were harvested and re-suspended in cell lysis buffer (50 mM Tris, 150 mM NaCl, 20 mM Imidazole, and 1% Triton X-100, pH 8.0). Before lysis, cOmplete™ Protease Inhibitor Cocktail (Sigma-Aldrich) and Dnase were added. Cells were lysed at 35 kPSI using a cell disruptor (Constant Systems) and the lysate centrifuged at 30,600 x *g* for 60 minutes. Clarified lysate was applied to gravity column containing Ni-NTA affinity resin pre-equilibrated with lysis buffer. The column was washed with 10 column volumes each of the following wash buffers: 1) 40 mM Tris-HCl, 300 mM NaCl, 1% Triton X-100, pH 8.0. 2) 40 mM Tris-HCl, 300 mM NaCl, 50 mM Sodium Cholate, 20 mM Imidazole, pH 8.0. 3) 40 mM Tris-HCl, 300 mM NaCl, 40 mM Imidazole, pH 8.0. His_6_-MSP2N2 was eluted with elution buffer (40 mM Tris-HCl, 300 mM NaCl, 400 mM Imidazole, pH 8.0). The eluted protein was dialyzed into 20 mM Tris-HCl, 100 mM NaCl, pH 8.0 buffer, concentrated and finally passed over a HiLoad 16/600 Superdex 200 pg column (Cytiva) using sizing buffer (40 mM Tris-HCl, 100 mM NaCl, pH 8.0) for size-exclusion chromatography (Extended Data Fig. 8c). SDS-PAGE analysis was used to estimate the purity of the sample (Extended Data Fig. 8d). Pure His_6_-MSP2N2 (≥99% w/w) was concentrated to 1.5 mg/ml, flash frozen in liquid N_2_ and stored at -70 °C until use.

### Nanodisc preparation

Lipid stocks dissolved in chloroform, purchased from Avanti® Polar Lipids, consisting of 18:1 PS (1,2-dioleoyl-sn-glycero-3-phospho-L-serine) and 14:0 CL (1’,3’-bis[1,2-dimyristoyl-sn-glycero-3-phospho]-glycerol) were mixed and dried under N2 gas to form dry lipid films. These were stored under vacuum overnight to ensure full removal of the solvent. Lipid films were solubilized in standard nanodisc buffer (20 mM Tris-HCl, 100 mM NaCl, 0.5 mM EDTA, pH 7.4) supplemented with 1.5% DDM and sonicated at 37 °C for 45 minutes. His_6_-MSP2N2 was incubated with DDM-solubilized lipids at a molar ratio of 1:11:98 (His_6_-MSP2N2:CL:PS) at 25 °C for 1 hour. Samples were dialyzed against standard nanodisc buffer (1:1000) for up to 30 hours (dialysis buffer was replaced at least 3 times). The sample was concentrated and run on a Superdex 200 Increase 10/300 GL using standard nanodisc buffer. Fractions containing His_6_-MSP2N2 were collected and incubated with endophilin-B1 at a molar ratio of 1:20 (His_6_-MSP2N2:EnB1) for 1 hour at 4 °C. The sample was concentrated and run on a Superdex 200 Increase 10/300 GL using sizing buffer (Extended Data Fig. 9a and 9b). Fractions containing both His_6_-MSP2N2 and endophilin-B1 were identified by western blot analysis using anti-EnB1 polyclonal (Goat; Invitrogen; Extended Data Fig. 9c) and anti-polyHis monoclonal primary antibodies (Mouse; Sigma Aldrich; Extended Data Fig. 9d) followed by alkaline phosphatase conjugated anti-goat or anti-mouse secondary antibodies (Sigma Aldrich). Membranes were stained using 1-Step™ NBT/BCIP Substrate Solution (Thermo Scientific).

### Cryo-EM sample preparation and data collection

SEC fractions containing endophilin-B1 and MSP2N2 were pooled and concentrated. Three microliters were applied to glow-discharged holey carbon grids (Quantifoil Micro Tools GmbH). Grids were blotted at 4°C and 95% humidity using filter paper (Whatman®) and plunge-frozen in liquid ethane using a Vitribot Mark IV (Thermo Fisher Scientific). Vitrified grids were screened at the Cryo-EM Uppsala facility using a 200 kV Glacios (Thermo Fisher Scientific) equipped with a Falcon 3EC direct electron detector (Thermo Fisher Scientific). Cryo-EM data were collected to confirm sample quality and produce initial 3D reconstructions. Endophilin B1-decorated nanodiscs data was collected at SciLifeLab using a Titan Krios G3i (Thermo Fisher Scientific) operated at 300 kV, equipped with a K3 BioQuantum direct electron detector (Gatan Inc.) and energy filter using 20 eV slit, at 130,000 x nominal magnification (Extended Data Table 1). Soluble endophilin B1 data was collected at the 3D-EM facility at Karolinska Institutet using a Titan Krios G3i operated at 300 kV, equipped with a cold-FEG (Thermo Fisher Scientific), a K3 BioQuantum direct electron detector (Gatan Inc.) and energy filter using 10 eV slit, at 165,000 x nominal magnification.

### Cryo-EM data processing

Movies from Glacios data collections were imported into cryoSPARC followed by patch motion correction, patch CTF correction and discarding of low-quality micrographs.^60^ Manually picked particles were filtered using 2D classification and 2D classes used for template picking. Junk particles were removed through multiple rounds of 2D classification followed by *ab-initio* reconstruction and non-uniform refinement.^88^

Movies from Krios data collections were imported, patch motion corrected and patch CTF corrected in cryoSPARC. Initial picking was performed using template picking and 2D classes generated using the final electron density map from the small-scale Glacios data collection on the same sample. Picked particles were filtered using the NCC score and then extracted using a box size of 512^2^ pixels. These particles were Fourier-cropped to 128^2^ pix for initial 2D classification and *ab-intio* reconstruction. Following the 2D classification and 3D reconstruction, the particles were re-extracted at 512^2^ pixels and Fourier-cropped to 256^2^ pix for further classification using several rounds of heterogeneous *ab-initio* and heterogeneous refinements. Final classes were refined using non-uniform refinement. Masks for individual endophilin B1 dimers were generated in UCSF ChimeraX and used in local refinements in cryoSPARC (Extended Data Table 1).^89^

### CryoDRGN

Particle sets were taken from different parts of the data processing “timeline” and analyzed with cryoDRGN.^61–63^ Before training, each particle set was Fourier-cropped to 128^2^ pixels in cryoDRGN. Larger and more heterogeneous data was processed over 30-60 epochs using the heterogeneous *ab-initio* reconstruction function with default architecture (3×256) and smaller less heterogeneous data was processed using the train_vae function, initially with the default architecture and later with a larger architecture (3×1024). Particle sets were split into groups based on kmeans filtering using the jupyter-notebook script provided in the cryoDRGN software package. Groups were separately re-imported to cryoSPARC for validation using *ab-initio* reconstruction and non-uniform refinement. Movies were created using a variation of the workflow described in ^90^ (https://github.com/Guillawme/).

### Model building

The AlphaFold model for endophilin B1 was processed in PHENIX.^52,53,91,92^ The model was rigid-body docked into the density maps in UCSF ChimeraX and refined in *Coot* and Servalcat.^89,93–95^ Model validation was performed using MolProbity (Extended Data Table 1).^96^

### Small-angle X-ray scattering measurements

All SAXS measurements were carried out on BM29 beamline (ESRF, Grenoble, France). All measurements were performed with 100% beam intensity at a wavelength of 0.9918 Å (12.5 keV). Initial data processing was performed automatically using the EDNA pipeline. See Extended Data Table 2 for other details of SAXS measurements.

### Small-angle scattering data processing

SAXS profiles (I(q)) were processed using ATSAS and BioXTAS RAW software suites.^97,98^ The protein concentrations were small, consequently the influence of structural factors on scattering curves was negligible. For calculation of values of molar absorption coefficient (ε), molecular mass, and scattering length density (SLD) from sequences, the programs ProtParam, Peptide Property Calculator and SLD calculator were used (see Extended Data Table 2). Distance distribution functions P(r) and regularized I(q) were obtained using GNOM program, which realizes the method of Indirect-Fourier Transform (IFT).^99^ Values of R_G_ and I(0) (Extended Data Table 2) were calculated from P(r) and using Guinier approximations. For model electron density fit and *ab-initio* electron density maps creation was used DENSS.^100^

### Liposome preparation

Lipid stocks dissolved in chloroform, 18:1 PS (1,2-dioleoyl-sn-glycero-3-phospho-L-serine), 18:1 PE (1,2-dioleoyl-sn-glycero-3-phosphoethanolamine), 18:1 CL (1’,3’-bis[1,2-dioleoyl-sn-glycero-3-phospho]-glycerol) and cholesterol, were mixed and dried under N_2_ gas to form dry lipid films. These were stored under vacuum overnight to ensure full removal of the solvent. Lipid films were solubilized in a 100 μM calcein (Merck) solution and extruded through a 1 μm filter. Calcein-encapsulated liposomes were separated from free calcein by gel filtration using G-50 Sephadex® resin (Cytiva) and sizing buffer. The encapsulation efficiency for each batch was determined by comparing liposomes alone (negative control) with liposomes in the presence of 1% Triton X-100 (Alfa Aesar), which fully permeabilizes liposomes. The fluorescence signal from positive controls were at least 10-fold higher than that of negative controls in all subsequent assays.

### Calcein release assay

Calcein encapsulated liposomes with different lipid compositions were added to endophilin B1 (final concentration 500 nM or 5 μM) in a 96-well plate and increase in calcein fluorescence was monitored in a CLARIOstar Plus plate reader (BMG Labtech) at excitation wavelength 482 nm and bandwidth 16 nm, emission wavelength 530 nm and bandwidth 40 nm with the dichroic mirror set to 504 nm. Sample fluorescence was determined every 5 minutes over a 60-minute period with shaking for 10 seconds prior to measuring. The results for each condition consist of at least 6 technical replicates. Negative controls consisted of calcein-encapsulated liposomes without protein. Positive controls consisted of calcein-encapsulated liposomes and 1% Triton X-100, which represented the maximum release of calcein. Negative and positive controls were included in each run and results were calculated using the following equation: %Calcein release = (*F_exp_ – F_neg_*)/(*F_pos_ – F_neg_*). Where *F_exp_* is the fluorescence of the sample, *F_0_* the average fluorescence of the negative controls and *F_pos_* the average signal from positive controls. The statistical relevance of results from different conditions was evaluated using paired t-tests.

## References

1. McMahon, H.T., and Gallop, J.L. (2005). Membrane curvature and mechanisms of dynamic cell membrane remodelling. Nature 438, 590–596. 10.1038/nature04396.

2. Peter, B.J., Kent, H.M., Mills, I.G., Vallis, Y., Butler, P.J.G., Evans, P.R., and McMahon, H.T. (2004). BAR Domains as Sensors of Membrane Curvature: The Amphiphysin BAR Structure. Science (1979) 303, 495– 499. 10.1126/SCIENCE.1092586.

3. Frost, A., Unger, V.M., and De Camilli, P. (2009). The BAR Domain Superfamily: Membrane-Molding Macromolecules. Cell 137, 191–196. 10.1016/J.CELL.2009.04.010.

4. Qualmann, B., Koch, D., and Kessels, M.M. (2011). Let’s go bananas: Revisiting the endocytic BAR code. EMBO Journal 30, 3501–3515. 10.1038/EMBOJ.2011.266.

5. Mim, C., and Unger, V.M. (2012). Membrane curvature and its generation by BAR proteins. Trends Biochem Sci 37, 526–533. 10.1016/J.TIBS.2012.09.001.

6. Simunovic, M., Voth, G.A., Callan-Jones, A., and Bassereau, P. (2015). When Physics Takes Over: BAR Proteins and Membrane Curvature. Trends Cell Biol 25, 780–792. 10.1016/J.TCB.2015.09.005.

7. Weissenhorn, W. (2005). Crystal structure of the endophilin-A1 BAR domain. J Mol Biol 351, 653–661. 10.1016/j.jmb.2005.06.013.

8. Gallop, J.L., Jao, C.C., Kent, H.M., Butler, P.J.G., Evans, P.R., Langen, R., and McMahon, H.T. (2006). Mechanism of endophilin N-BAR domain-mediated membrane curvature. EMBO Journal 25, 2898–2910. 10.1038/sj.emboj.7601174.

9. Takei, K., Slepnev, V.I., Haucke, V., and De Camilli, P.(1999). Functional partnership between amphiphysin and dynamin in clathrin-mediated endocytosis.

10. Masuda, M., Takeda, S., Sone, M., Ohki, T., Mori, H., Kamioka, Y., and Mochizuki, N. (2006). Endophilin BAR domain drives membrane curvature by two newly identified structure-based mechanisms. EMBO Journal 25, 2889–2897. 10.1038/sj.emboj.7601176.

11. Suetsugu, S., Murayama, K., Sakamoto, A., Hanawa-Suetsugu, K., Seto, A., Oikawa, T., Mishima, C., Shirouzu, M., Takenawa, T., and Yokoyama, S. (2006). The RAC binding domain/IRSp53-MIM homology domain of IRSp53 induces RAC-dependent membrane deformation. Journal of Biological Chemistry 281, 35347–35358. 10.1074/jbc.M606814200.

12. Henne, W.M., Kent, H.M., Ford, M.G.J., Hegde, B.G., Daumke, O., Butler, P.J.G., Mittal, R., Langen, R., Evans, P.R., and McMahon, H.T. (2007). Structure and Analysis of FCHo2 F-BAR Domain: A Dimerizing and Membrane Recruitment Module that Effects Membrane Curvature. Structure 15, 839–852. 10.1016/j.str.2007.05.002.

13. Mattila, P.K., Pykäläinen, A., Saarikangas, J., Paavilainen, V.O., Vihinen, H., Jokitalo, E., and Lappalainen, P. (2007). Missing-in-metastasis and IRSp53 deform PI(4,5)P2-rich membranes by an inverse BAR domain-like mechanism. Journal of Cell Biology 176, 953–964. 10.1083/jcb.200609176.

14. Pylypenko, O., Lundmark, R., Rasmuson, E., Carlsson, S.R., and Rak, A. (2007). The PX-BAR membrane-remodeling unit of sorting nexin 9. EMBO Journal 26, 4788–4800. 10.1038/sj.emboj.7601889.

15. Shimada, A., Niwa, H., Tsujita, K., Suetsugu, S., Nitta, K., Hanawa-Suetsugu, K., Akasaka, R., Nishino, Y., Toyama, M., Chen, L., et al. (2007). Curved EFC/F-BAR-Domain Dimers Are Joined End to End into a Filament for Membrane Invagination in Endocytosis. Cell 129, 761–772. 10.1016/j.cell.2007.03.040.

16. Frost, A., Perera, R., Roux, A., Spasov, K., Destaing, O., Egelman, E.H., De Camilli, P., and Unger, V.M. (2008). Structural Basis of Membrane Invagination by F-BAR Domains. Cell 132, 807–817. 10.1016/j.cell.2007.12.041.

17. Reider, A., Barker, S.L., Mishra, S.K., Im, Y.J., Maldonado-Báez, L., Hurley, J.H., Traub, L.M., and Wendland, B. (2009). Syp1 is a conserved endocytic adaptor that contains domains involved in cargo selection and membrane tubulation. EMBO Journal 28, 3103–3116. 10.1038/emboj.2009.248.

18. Saarikangas, J., Zhao, H., Pykäläinen, A., Laurinmäki, P., Mattila, P.K., Kinnunen, P.K.J., Butcher, S.J., and Lappalainen, P. (2009). Molecular Mechanisms of Membrane Deformation by I-BAR Domain Proteins. Current Biology 19, 95–107. 10.1016/j.cub.2008.12.029.

19. Wang, Q., Navarro, M.V.A.S., Peng, G., Molinelli, E., Lin Goh, S., Judson, B.L., Rajashankar, K.R., and Sondermann, H. (2009). Molecular mechanism of membrane constriction and tubulation mediated by the F-BAR protein Pacsin/Syndapin. Proceedings of the National Academy of Sciences 106, 12700–12705. 10.1073/pnas.0902974106.

20. Rao, Y., Ma, Q., Vahedi-Faridi, A., Sundborger, A., Pechstein, A., Puchkov, D., Luo, L., Shupliakov, O., Saenger, W., and Haucke, V. (2010). Molecular basis for SH3 domain regulation of F-BAR-mediated membrane deformation. Proc Natl Acad Sci U S A 107, 8213–8218. 10.1073/PNAS.1003478107/SUPPL_FILE/PNAS.201003478SI.PDF.

21. Shimada, A., Takano, K., Shirouzu, M., Hanawa-Suetsugu, K., Terada, T., Toyooka, K., Umehara, T., Yamamoto, M., Yokoyama, S., and Suetsugu, S. (2010). Mapping of the basic amino-acid residues responsible for tubulation and cellular protrusion by the EFC/F-BAR domain of pacsin2/Syndapin II. FEBS Lett 584, 1111–1118. 10.1016/j.febslet.2010.02.058.

22. Sundborger, A., Soderblom, C., Vorontsova, O., Evergren, E., Hinshaw, J.E., and Shupliakov, O. (2011). An endophilin-dynamin complex promotes budding of clathrin-coated vesicles during synaptic vesicle recycling. J Cell Sci 124, 133–143. 10.1242/jcs.072686.

23. Mim, C., Cui, H., Gawronski-Salerno, J.A., Frost, A., Lyman, E., Voth, G.A., and Unger, V.M. (2012). Structural basis of membrane bending by the N-BAR protein endophilin. Cell 149, 137–145. 10.1016/j.cell.2012.01.048.

24. Pang, X., Fan, J., Zhang, Y., Zhang, K., Gao, B., Ma, J., Li, J., Deng, Y., Zhou, Q., Egelman, E.H., et al. (2014). A PH domain in ACAP1 possesses key features of the BAR domain in promoting membrane curvature. Dev Cell 31, 73–86. 10.1016/j.devcel.2014.08.020.

25. Bhatt, V.S., Ashley, R., and Sundborger-Lunna, A. (2021). Amphipathic Motifs Regulate N-BAR Protein Endophilin B1 Auto-inhibition and Drive Membrane Remodeling. Structure 29, 61–69.e3. 10.1016/j.str.2020.09.012.

26. Lopez-Robles, C., Scaramuzza, S., Astorga-Simon, E.N., Ishida, M., Williamson, C.D., Baños-Mateos, S., Gil-Carton, D., Romero-Durana, M., Vidaurrazaga, A., Fernandez-Recio, J., et al. (2023). Architecture of the ESCPE-1 membrane coat. Nat Struct Mol Biol 30, 958–969. 10.1038/s41594-023-01014-7.

27. Etxebarria, A., Terrones, O., Yamaguchi, H., Landajuela, A., Landeta, O., Antonsson, B., Wang, H.G., and Basañez, G. (2009). Endophilin B1/Bif-1 stimulates BAX activation independently from its capacity to produce large scale membrane morphological rearrangements. Journal of Biological Chemistry 284, 4200– 4212. 10.1074/jbc.M808050200.

28. Karbowski, M., Jeong, S.Y., and Youle, R.J. (2004). Endophilin B1 is required for the maintenance of mitochondrial morphology. Journal of Cell Biology 166, 1027–1039. 10.1083/jcb.200407046.

29. Takahashi, Y., Karbowski, M., Yamaguchi, H., Kazi, A., Wu, J., Sebti, S.M., Youle, R.J., and Wang, H.-G. (2005). Loss of Bif-1 Suppresses Bax/Bak Conformational Change and Mitochondrial Apoptosis. Mol Cell Biol 25, 9369–9382. 10.1128/mcb.25.21.9369-9382.2005.

30. Pierrat, B., Simonen, M., Cueto, M., Mestan, J., Ferrigno, P., and Heim, J. (2001). SH3GLB, a New Endophilin-Related Protein Family Featuring an SH3 Domain. Genomics 71, 222–234. 10.1006/geno.2000.6378.

31. Cuddeback, S.M., Yamaguchi, H., Komatsu, K., Miyashita, T., Yamada, M., Wu, C., Singh, S., and Wang, H.G. (2001). Molecular cloning and characterization of Bif-1. A novel Src homology 3 domain-containing protein that associates with Bax. Journal of Biological Chemistry 276, 20559–20565. 10.1074/jbc.M101527200.

32. Robustelli, J., and Baumgart, T. (2021). Membrane partitioning and lipid selectivity of the N-terminal amphipathic H0 helices of endophilin isoforms. Biochim Biophys Acta Biomembr 1863. 10.1016/j.bbamem.2021.183660.

33. Takahashi, Y., Coppola, D., Matsushita, N., Cualing, H.D., Sun, M., Sato, Y., Liang, C., Jung, J.U., Cheng, J.Q., Mulé, J.J., et al. (2007). Bif-1 interacts with Beclin 1 through UVRAG and regulates autophagy and tumorigenesis. Nat Cell Biol 9, 1142–1151. 10.1038/NCB1634.

34. Thoresen, S.B., Pedersen, N.M., Liestøl, K., and Stenmark, H. (2010). A phosphatidylinositol 3-kinase class III sub-complex containing VPS15, VPS34, Beclin 1, UVRAG and BIF-1 regulates cytokinesis and degradative endocytic traffic. Exp Cell Res 316, 3368–3378. 10.1016/j.yexcr.2010.07.008.

35. Wang, Y.H., Wang, J.Q., Wang, Q., Wang, Y., Guo, C., Chen, Q., Chai, T., and Tang, T.S. (2016). Endophilin B2 promotes inner mitochondrial membrane degradation by forming heterodimers with Endophilin B1 during mitophagy. Sci Rep 6. 10.1038/srep25153.

36. Cho, S.G., Xiao, X., Wang, S., Gao, H., Rafikov, R., Black, S., Huang, S., Ding, H.F., Yoon, Y., Kirken, R.A., et al. (2019). Bif-1 interacts with prohibitin-2 to regulate mitochondrial inner membrane during cell stress and apoptosis. Journal of the American Society of Nephrology 30, 1174–1191. 10.1681/ASN.2018111117.

37. Bonner, A.E., Lemon, W.J., Devereux, T.R., Lubet, R.A., and You, M. (2004). Molecular profiling of mouse lung tumors: association with tumor progression, lung development, and human lung adenocarcinomas. Oncogene 23, 1166–1176. 10.1038/sj.onc.1207234.

38. Coppola, D., Khalil, F., Eschrich, S.A., Boulware, D., Yeatman, T., and Wang, H. (2008). Down-regulation of Bax-interacting factor-1 in colorectal adenocarcinoma. Cancer 113, 2665–2670. 10.1002/cncr.23892.

39. Kim, S.Y., Oh, Y.L., Kim, K.M., Jeong, E.G., Kim, M.S., Yoo, N.J., and Lee, S.H. (2008). Decreased expression of Bax-interacting factor-1 (Bif-1) in invasive urinary bladder and gallbladder cancers. Pathology 40, 553–557. 10.1080/00313020802320440.

40. Ho, J., Kong, J.-W.-F., Choong, L.-Y., Loh, M.-C.-S., Toy, W., Chong, P.-K., Wong, C.-H., Wong, C.-Y., Shah, N., and Lim, Y.-P. (2009). Novel Breast Cancer Metastasis-Associated Proteins. J Proteome Res 8, 583–594. 10.1021/pr8007368.

41. Coppola, D., Helm, J., Ghayouri, M., Malafa, M.P., and Wang, H.-G. (2011). Down-Regulation of Bax-Interacting Factor 1 in Human Pancreatic Ductal Adenocarcinoma. Pancreas 40, 433–437. 10.1097/MPA.0b013e318205eb03.

42. Runkle, K.B., Meyerkord, C.L., Desai, N. V., Takahashi, Y., and Wang, H.-G. (2012). Bif-1 suppresses breast cancer cell migration by promoting EGFR endocytic degradation. Cancer Biol Ther 13, 956–966. 10.4161/cbt.20951.

43. Ko, Y.H., Cho, Y.-S., Won, H.S., An, H.J., Sun, D.S., Hong, S.U., Park, J.H., and Lee, M.A. (2013). Stage-Stratified Analysis of Prognostic Significance of Bax-Interacting Factor-1 Expression in Resected Colorectal Cancer. Biomed Res Int 2013, 1–8. 10.1155/2013/329839.

44. Xu, L., Wang, Z., He, S.-Y., Zhang, S.-F., Luo, H.-J., Zhou, K., Li, X.-F., Qiu, S.-P., and Cao, K.-Y. (2016). Bax-interacting factor-1 inhibits cell proliferation and promotes apoptosis in prostate cancer cells. Oncol Rep 36, 3513–3521. 10.3892/or.2016.5172.

45. Peter, B.J., Kent, H.M., Mills, I.G., Vallis, Y., Butler, P.J.G., Evans, P.R., and McMahon, H.T. (2004). BAR Domains as Sensors of Membrane Curvature: The Amphiphysin BAR Structure. Science (1979) 303, 495– 499. 10.1126/science.1092586.

46. Trempe, J.-F., Chen, C.X.-Q., Grenier, K., Camacho, E.M., Kozlov, G., McPherson, P.S., Gehring, K., and Fon, E.A. (2009). SH3 Domains from a Subset of BAR Proteins Define a Ubl-Binding Domain and Implicate Parkin in Synaptic Ubiquitination. Mol Cell 36, 1034–1047. 10.1016/j.molcel.2009.11.021.

47. Bai, X., Meng, G., Luo, M., and Zheng, X. (2012). Rigidity of wedge loop in PACSIN 3 protein is a key factor in dictating diameters of tubules. Journal of Biological Chemistry 287, 22387–22396. 10.1074/jbc.M112.358960.

48. Löw, C., Weininger, U., Lee, H., Schweimer, K., Neundorf, I., Beck-Sickinger, A.G., Pastor, R.W., and Balbach, J. (2008). Structure and dynamics of helix-0 of the N-BAR domain in lipid micelles and bilayers. Biophys J 95, 4315–4323. 10.1529/biophysj.108.134155.

49. Mizuno, N., Jao, C.C., Langen, R., and Steven, A.C. (2010). Multiple modes of endophilin-mediated conversion of lipid vesicles into coated tubes: Implications for synaptic endocytosis. Journal of Biological Chemistry 285, 23351–23358. 10.1074/jbc.M110.143776.

50. Kovtun, O., Leneva, N., Bykov, Y.S., Ariotti, N., Teasdale, R.D., Schaffer, M., Engel, B.D., Owen, D.J., Briggs, J.A.G., and Collins, B.M. (2018). Structure of the membrane-assembled retromer coat determined by cryo-electron tomography. Nature 561, 561–564. 10.1038/s41586-018-0526-z.

51. Allerston, C.K., Krojer, T., Cooper, C.D.O., Vollmar, M., Arrowsmith, C.H., Edwards, A., Bountra, C., von Delft, F., and Gileadi, O. (2012). Crystal structure of the N-BAR domain of human bridging integrator 2. 10.2210/pdb4avm/pdb.

52. Jumper, J., Evans, R., Pritzel, A., Green, T., Figurnov, M., Ronneberger, O., Tunyasuvunakool, K., Bates, R., Žídek, A., Potapenko, A., et al. (2021). Highly accurate protein structure prediction with AlphaFold. Nature 596, 583–589. 10.1038/s41586-021-03819-2.

53. Varadi, M., Anyango, S., Deshpande, M., Nair, S., Natassia, C., Yordanova, G., Yuan, D., Stroe, O., Wood, G., Laydon, A., et al. (2022). AlphaFold Protein Structure Database: Massively expanding the structural coverage of protein-sequence space with high-accuracy models. Nucleic Acids Res 50, D439–D444. 10.1093/nar/gkab1061.

54. Bayburt, T.H., and Sligar, S.G. (2002). Single-molecule height measurements on microsomal cytochrome P450 in nanometer-scale phospholipid bilayer disks. Proceedings of the National Academy of Sciences 99, 6725–6730. 10.1073/pnas.062565599.

55. Ritchie, T.K., Grinkova, Y.V., Bayburt, T.H., Denisov, I.G., Zolnerciks, J.K., Atkins, W.M., and Sligar, S.G. (2009). Reconstitution of Membrane Proteins in Phospholipid Bilayer Nanodiscs. In, pp. 211–231. 10.1016/S0076-6879(09)64011-8.

56. Denisov, I.G., and Sligar, S.G. (2016). Nanodiscs for structural and functional studies of membrane proteins. Nat Struct Mol Biol 23, 481–486. 10.1038/nsmb.3195.

57. Mclean, M.A., Gregory, M.C., and Sligar, S.G. (2018). Nanodiscs: A Controlled Bilayer Surface for the Study of Membrane Proteins. 10.1146/annurev-biophys.

58. Sligar, S.G., and Denisov, I.G. (2021). Nanodiscs: A toolkit for membrane protein science. Preprint at Blackwell Publishing Ltd, 10.1002/pro.3994.

59. S. Cannon, K., Sarsam, R.D., Tedamrongwanish, T., Zhang, K., and Baker, R.W. (2023). Lipid nanodiscs as a template for high-resolution cryo-EM structures of peripheral membrane proteins. J Struct Biol 215. 10.1016/j.jsb.2023.107989.

60. Punjani, A., Rubinstein, J.L., Fleet, D.J., and Brubaker, M.A. (2017). cryoSPARC: algorithms for rapid unsupervised cryo-EM structure determination. Nat Methods 14, 290–296. 10.1038/nmeth.4169.

61. Zhong, E.D., Bepler, T., Berger, B., and Davis, J.H. (2021). CryoDRGN: reconstruction of heterogeneous cryo-EM structures using neural networks. Nat Methods 18, 176–185. 10.1038/s41592-020-01049-4.

62. Kinman, L.F., Powell, B.M., Zhong, E.D., Berger, B., and Davis, J.H. (2022). Uncovering structural ensembles from single-particle cryo-EM data using cryoDRGN. Nature Protocols 2022 18:2 18, 319–339. 10.1038/S41596-022-00763-X.

63. Zhong, E.D., Lerer, A., Davis, J.H., and Berger, B. (2021). CryoDRGN2: Ab Initio Neural Reconstruction of 3D Protein Structures From Real Cryo-EM Images. In Proceedings of the IEEE/CVF International Conference on Computer Vision (ICCV), pp. 4066–4075.

64. Wang, Q., Kaan, H.Y.K., Hooda, R.N., Goh, S.L., and Sondermann, H. (2008). Structure and Plasticity of Endophilin and Sorting Nexin 9. Structure 16, 1574–1587. 10.1016/j.str.2008.07.016.

65. Shai, Y. (1999). Mechanism of the binding, insertion and destabilization of phospholipid bilayer membranes by α-helical antimicrobial and cell non-selective membrane-lytic peptides. Biochimica et Biophysica Acta (BBA) - Biomembranes 1462, 55–70. 10.1016/S0005-2736(99)00200-X.

66. Tosatto, L., Andrighetti, A.O., Plotegher, N., Antonini, V., Tessari, I., Ricci, L., Bubacco, L., and Dalla Serra, M. (2012). Alpha-synuclein pore forming activity upon membrane association. Biochimica et Biophysica Acta (BBA) - Biomembranes 1818, 2876–2883. 10.1016/j.bbamem.2012.07.007.

67. Boucrot, E., Pick, A., Çamdere, G., Liska, N., Evergren, E., McMahon, H.T., and Kozlov, M.M. (2012). Membrane fission is promoted by insertion of amphipathic helices and is restricted by crescent BAR domains. Cell 149, 124–136. 10.1016/j.cell.2012.01.047.

68. Stefanovic, A.N.D., Stöckl, M.T., Claessens, M.M.A.E., and Subramaniam, V. (2014). α-Synuclein oligomers distinctively permeabilize complex model membranes. FEBS J 281, 2838–2850. 10.1111/febs.12824.

69. Guha, S., Ghimire, J., Wu, E., and Wimley, W.C. (2019). Mechanistic Landscape of Membrane-Permeabilizing Peptides. Chem Rev 119, 6040–6085. 10.1021/acs.chemrev.8b00520.

70. Basu Ball, W., Neff, J.K., and Gohil, V.M. (2018). The role of nonbilayer phospholipids in mitochondrial structure and function. FEBS Lett 592, 1273–1290. 10.1002/1873-3468.12887.

71. Gasanov, S.E., Kim, A.A., Yaguzhinsky, L.S., and Dagda, R.K. (2018). Non-bilayer structures in mitochondrial membranes regulate ATP synthase activity. Biochimica et Biophysica Acta (BBA) - Biomembranes 1860, 586–599. 10.1016/j.bbamem.2017.11.014.

72. Imura, T., Tsukui, Y., Taira, T., Aburai, K., Sakai, K., Sakai, H., Abe, M., and Kitamoto, D. (2014). Surfactant-like properties of an amphiphilic α-helical peptide leading to lipid nanodisc formation. Langmuir 30, 4752– 4759. 10.1021/la500267b.

73. Frauenfeld, J., Löving, R., Armache, J.P., Sonnen, A.F.P., Guettou, F., Moberg, P., Zhu, L., Jegerschöld, C., Flayhan, A., Briggs, J.A.G., et al. (2016). A saposin-lipoprotein nanoparticle system for membrane proteins. Nat Methods 13, 345–351. 10.1038/nmeth.3801.

74. Bulankina, A. V., Deggerich, A., Wenzel, D., Mutenda, K., Wittmann, J.G., Rudolph, M.G., Burger, K.N.J., and Höning, S. (2009). TIP47 functions in the biogenesis of lipid droplets. Journal of Cell Biology 185, 641–655. 10.1083/jcb.200812042.

75. Varkey, J., Mizuno, N., Hegde, B.G., Cheng, N., Steven, A.C., and Langen, R. (2013). α-synuclein oligomers with broken helical conformation form lipoprotein nanoparticles. Journal of Biological Chemistry 288, 17620–17630. 10.1074/jbc.M113.476697.

76. Simunovic, M., Manneville, J.B., Renard, H.F., Evergren, E., Raghunathan, K., Bhatia, D., Kenworthy, A.K., Voth, G.A., Prost, J., McMahon, H.T., et al. (2017). Friction Mediates Scission of Tubular Membranes Scaffolded by BAR Proteins. Cell 170, 172–184.e11. 10.1016/j.cell.2017.05.047.

77. Aryal, C.M., Bui, N.N., Khadka, N.K., Song, L., and Pan, J. (2020). The helix 0 of endophilin modifies membrane material properties and induces local curvature. Biochim Biophys Acta Biomembr 1862. 10.1016/j.bbamem.2020.183397.

78. Ringstad, N., Gad, H., Löw, P., Di Paolo, G., Brodin, L., Shupliakov, O., and De Camilli, P. (1999). Endophilin/SH3p4 Is Required for the Transition from Early to Late Stages in Clathrin-Mediated Synaptic Vesicle Endocytosis. Neuron 24, 143–154. 10.1016/S0896-6273(00)80828-4.

79. Guichet, A., Wucherpfennig, T., Dudu, V., Etter, S., Wilsch-Bräuniger, M., Hellwig, A., González-Gaitán, M., Huttner, W.B., and Schmidt, A.A. (2002). Essential role of endophilin A in synaptic vesicle budding at the Drosophila neuromuscular junction. EMBO J 21, 1661–1672. 10.1093/emboj/21.7.1661.

80. Hovius, R., Lambrechts, H., Nicolay, K., and de Kruijff, B. (1990). Improved methods to isolate and subfractionate rat liver mitochondria. Lipid composition of the inner and outer membrane. Biochimica et Biophysica Acta (BBA) - Biomembranes 1021, 217–226. 10.1016/0005-2736(90)90036-N.

81. Holthuis, J.C.M., and Menon, A.K. (2014). Lipid landscapes and pipelines in membrane homeostasis. Nature 510, 48–57. 10.1038/nature13474.

82. Kagan, V.E., Tyurin, V.A., Jiang, J., Tyurina, Y.Y., Ritov, V.B., Amoscato, A.A., Osipov, A.N., Belikova, N.A., Kapralov, A.A., Kini, V., et al. (2005). Cytochrome c acts as a cardiolipin oxygenase required for release of proapoptotic factors. Nat Chem Biol 1, 223–232. 10.1038/nchembio727.

83. Belikova, N.A., Vladimirov, Y.A., Osipov, A.N., Kapralov, A.A., Tyurin, V.A., Potapovich, M. V., Basova, L. V., Peterson, J., Kurnikov, I. V., and Kagan, V.E. (2006). Peroxidase Activity and Structural Transitions of Cytochrome *c* Bound to Cardiolipin-Containing Membranes. Biochemistry 45, 4998–5009. 10.1021/bi0525573.

84. Chu, C.T., Ji, J., Dagda, R.K., Jiang, J.F., Tyurina, Y.Y., Kapralov, A.A., Tyurin, V.A., Yanamala, N., Shrivastava, I.H., Mohammadyani, D., et al. (2013). Cardiolipin externalization to the outer mitochondrial membrane acts as an elimination signal for mitophagy in neuronal cells. Nat Cell Biol 15, 1197–1205. 10.1038/ncb2837.

85. Schug, Z.T., and Gottlieb, E. (2009). Cardiolipin acts as a mitochondrial signalling platform to launch apoptosis. Biochimica et Biophysica Acta (BBA) - Biomembranes 1788, 2022–2031. 10.1016/j.bbamem.2009.05.004.

86. Grinkova, Y. V., Denisov, I.G., and Sligar, S.G. (2010). Engineering extended membrane scaffold proteins for self-assembly of soluble nanoscale lipid bilayers. Protein Engineering Design and Selection 23, 843–848. 10.1093/protein/gzq060.

87. Bayburt, T.H., Grinkova, Y. V., and Sligar, S.G. (2002). Self-Assembly of Discoidal Phospholipid Bilayer Nanoparticles with Membrane Scaffold Proteins. Nano Lett 2, 853–856. 10.1021/nl025623k.

88. Punjani, A., Zhang, H., and Fleet, D.J. (2020). Non-uniform refinement: adaptive regularization improves single-particle cryo-EM reconstruction. Nat Methods 17, 1214–1221. 10.1038/s41592-020-00990-8.

89. Pettersen, E.F., Goddard, T.D., Huang, C.C., Meng, E.C., Couch, G.S., Croll, T.I., Morris, J.H., and Ferrin, T.E. (2021). UCSF ChimeraX: structure visualization for researchers, educators, and developers. Protein Sci. 30, 70–82. 10.1002/pro.3943.

90. Bacic, L., Gaullier, G., Sabantsev, A., Lehmann, L.C., Brackmann, K., Dimakou, D., Halic, M., Hewitt, G., Boulton, S.J., and Deindl, S. (2021). Structure and dynamics of the chromatin remodeler ALC1 bound to a PARylated nucleosome. Elife 10. 10.7554/eLife.71420.

91. Liebschner, D., Afonine, P. V., Baker, M.L., Bunkoczi, G., Chen, V.B., Croll, T.I., Hintze, B., Hung, L.W., Jain, S., McCoy, A.J., et al. (2019). Macromolecular structure determination using X-rays, neutrons and electrons: Recent developments in Phenix. Acta Crystallogr D Struct Biol 75, 861–877. 10.1107/S2059798319011471/HTTPS://GOLDBOOK.IUPAC.ORG/E02002.HTML.

92. Hiranuma, N., Park, H., Baek, M., Anishchenko, I., Dauparas, J., and Baker, D. (2021). Improved protein structure refinement guided by deep learning based accuracy estimation. Nat Commun 12. 10.1038/s41467-021-21511-x.

93. Emsley, P., Lohkamp, B., Scott, W.G., and Cowtan, K. (2010). Features and development of Coot. Acta Crystallogr. Sect. D 66, 486–501. 10.1107/s0907444910007493.

94. Casañal, A., Lohkamp, B., and Emsley, P. (2020). Current developments in Coot for macromolecular model building of Electron Cryo-microscopy and Crystallographic Data. Protein Science 29, 1069–1078. 10.1002/PRO.3791.

95. Yamashita, K., Palmer, C.M., Burnley, T., and Murshudov, G.N. (2021). Cryo-EM single-particle structure refinement and map calculation using Servalcat. Acta Crystallogr D Struct Biol 77, 1282–1291. 10.1107/S2059798321009475.

96. Chen, V.B., Arendall, W.B., Headd, J.J., Keedy, D.A., Immormino, R.M., Kapral, G.J., Murray, L.W., Richardson, J.S., and Richardson, D.C. (2010). MolProbity: All-atom structure validation for macromolecular crystallography. Acta Crystallogr D Biol Crystallogr 66, 12–21. 10.1107/S0907444909042073.

97. Hopkins, J.B., Gillilan, R.E., and Skou, S. (2017). *BioXTAS RAW* : improvements to a free open-source program for small-angle X-ray scattering data reduction and analysis. J Appl Crystallogr 50, 1545–1553. 10.1107/S1600576717011438.

98. Manalastas-Cantos, K., Konarev, P. V., Hajizadeh, N.R., Kikhney, A.G., Petoukhov, M. V., Molodenskiy, D.S., Panjkovich, A., Mertens, H.D.T., Gruzinov, A., Borges, C., et al. (2021). *ATSAS 3.0* : expanded functionality and new tools for small-angle scattering data analysis. J Appl Crystallogr 54, 343–355. 10.1107/S1600576720013412.

99. Semenyuk, A. V., and Svergun, D.I. (1991). GNOM – a program package for small-angle scattering data processing. J Appl Crystallogr 24, 537–540. 10.1107/S002188989100081X.

100. Grant, T.D. (2018). Ab initio electron density determination directly from solution scattering data. Nat Methods 15, 191–193. 10.1038/nmeth.4581.

