## Extended data for "Peripheral membrane protein endophilin B1 probes, perturbs and permeabilizes lipid bilayers"

### Extended Data Figures

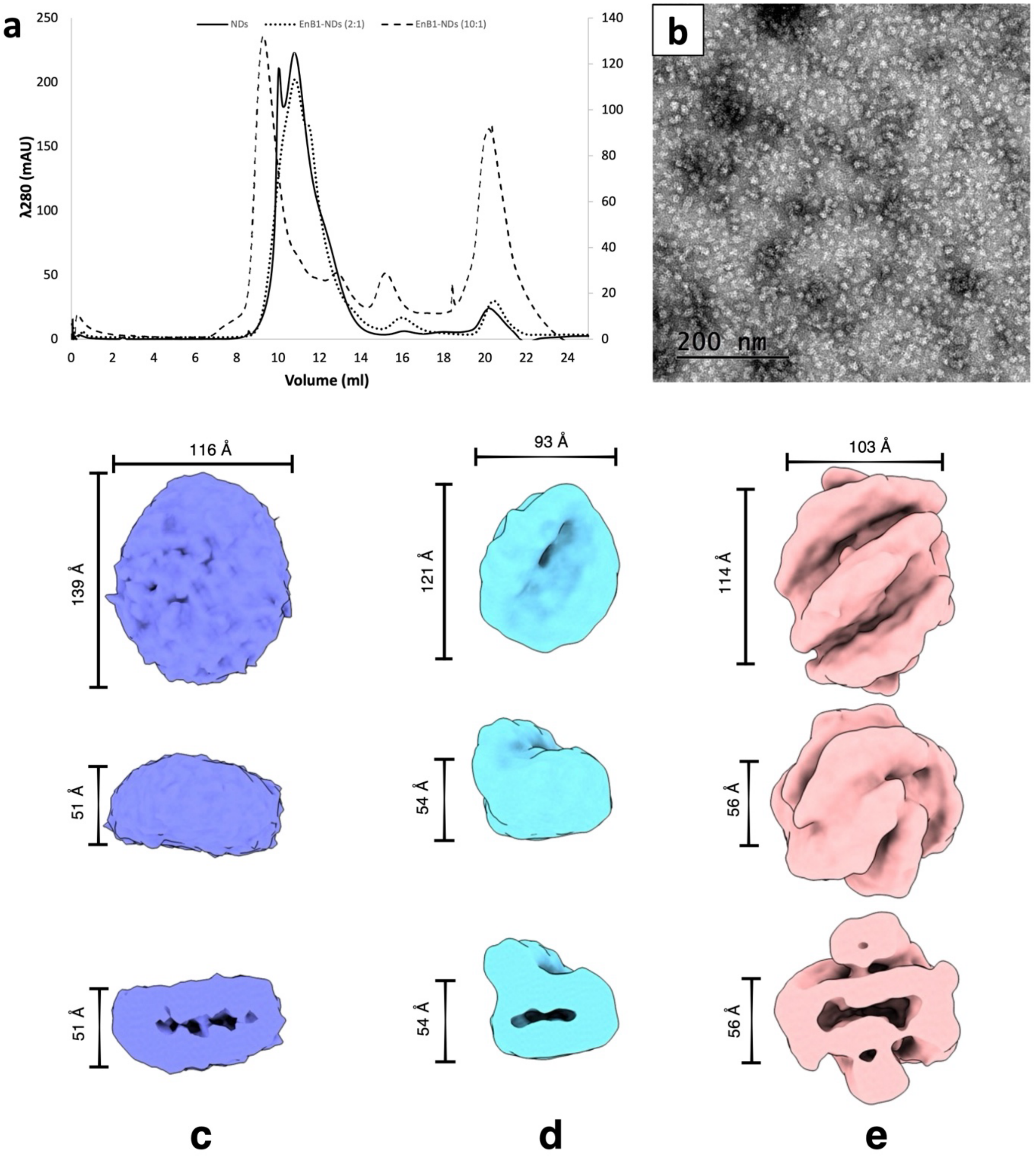

Extended Data Figure 1. Higher ratios of endophilin B1 (EnB1) yield nanodiscs (NDs) with more EnB1 decoration. **a**) SEC from Superdex 200 Increase 10/300 analysis of NDs, EnB1+NDs (1:2 of MSP2N2: EnB1), and EnB1+NDs (1:10 of MSP2N2: EnB1). **b**) Negative stain micrograph showing the EnB1 10:1 peak that eluted at 9.3 ml visualized by 1% uranyl acetate. Cryo-EM reconstructions of naked MSP2N2 NDs (**c**), EnB1 ND complex (1:2 of MSP2N2:EnB1) (**d**), and EnB1 ND complex (1:10 of MSP2N2:EnB1) (**e**) from Glacios data sets. EnB1 binding reduces the diameter of MSP2N2 NDs by roughly 20 Å and the binding of 6 EnB1 dimers changes the shape of the disc.

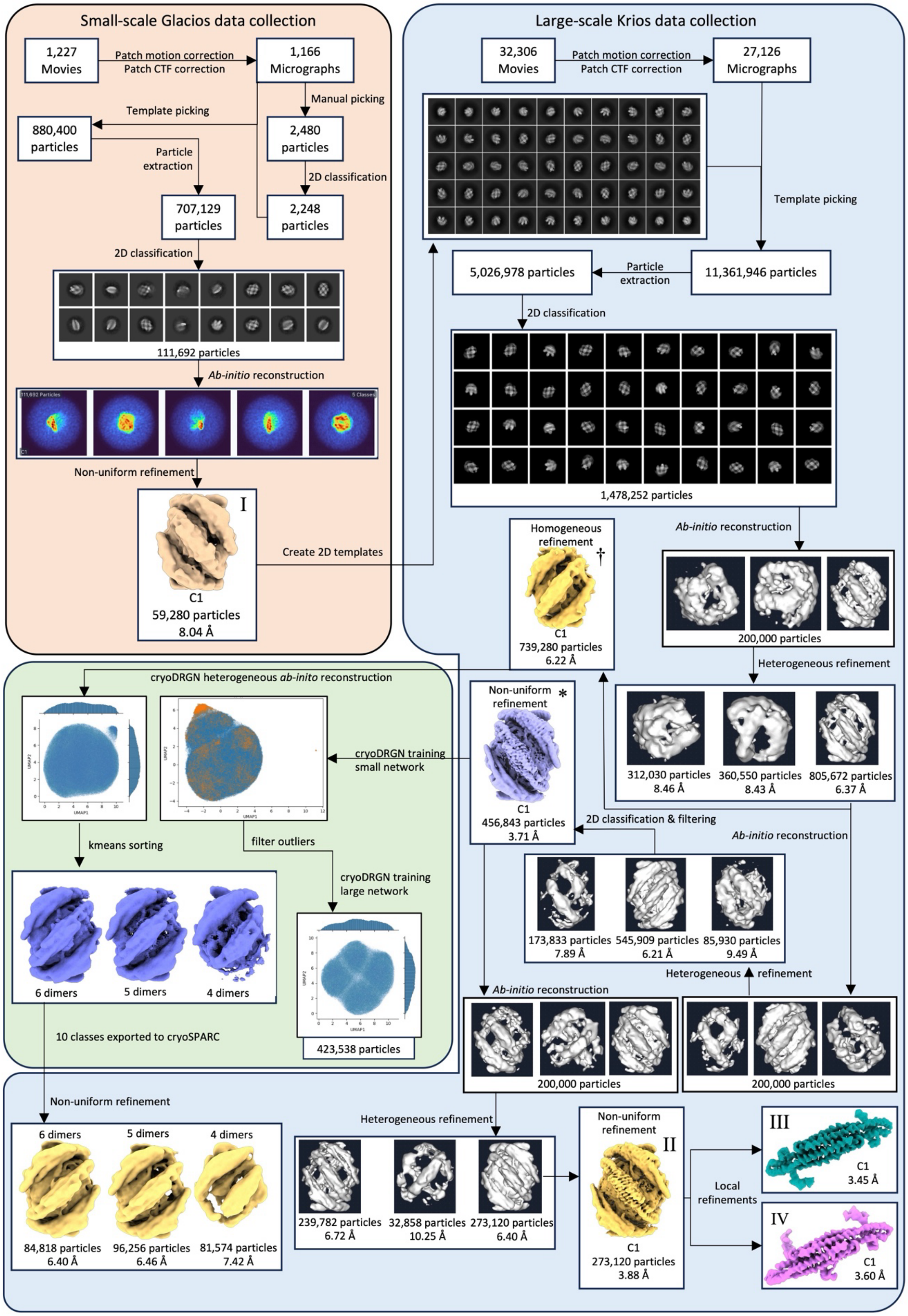

Extended data figure 2. Cryo-EM workflow summary.

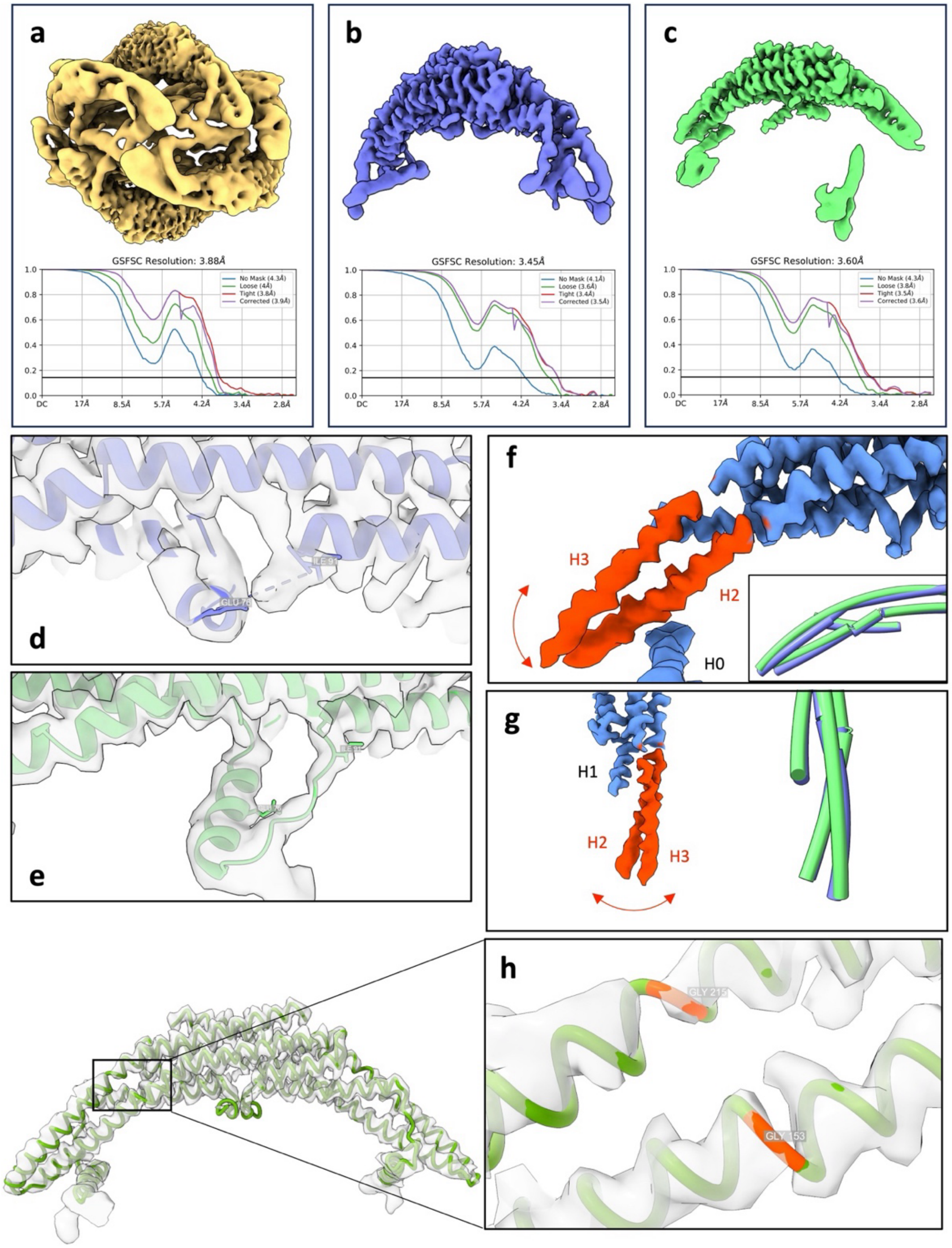

Extended data figure 3. Cryo-EM maps and subsequent models reveal distinct conformations of amphipathic regions and the BAR domain. **a**) Final C1 consensus map and gold-standard Fourier-shell correlation (GSFSC) curve. **b**) Center dimer focused map and GSFSC curve. **c**) Side dimer focused map and GSFSC curve. **d**) Center dimer map and atomic model highlighting that H1i is disordered in solution. **e**) Side dimer map and atomic model highlighting that H1i assumes a mostly helical conformation when membrane-inserted **f-g**) The distal ends of H2 and H3 helices (red) have worse local resolution due to flexibility around the hinge region. The position of these is also different in the atomic models (center dimer in blue and side dimer in green). **h**) Center dimer map and model highlighting the two glycines, Gly153 and Gly215, that are responsible for the increased flexibility of the distal ends of the BAR domain.

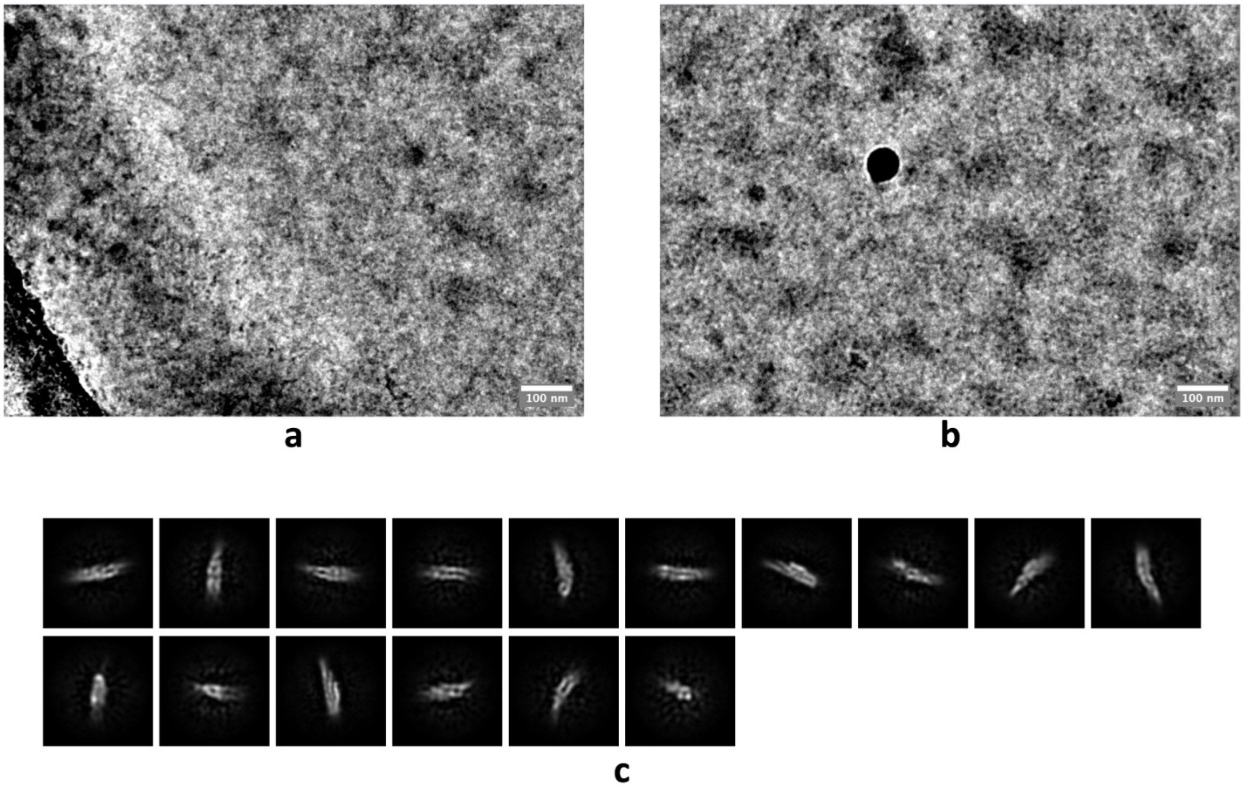

Extended data figure 4. Cryo-EM of *wt* EnB1 in solution. **a-b**) Raw micrographs from high-resolution data collection of soluble EnB1, low-pass filtered to 8 Å. **c**) 2D classes of soluble EnB1 (43,341 particles).

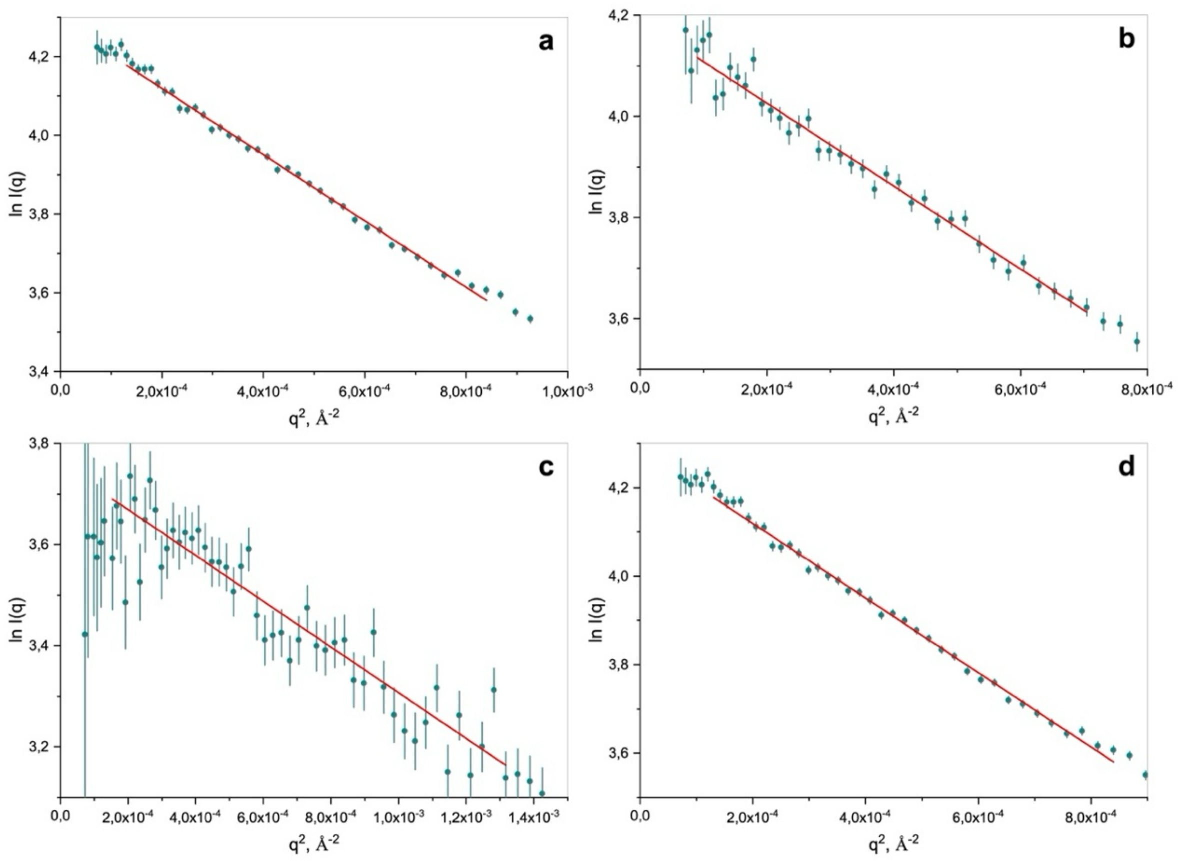

Extended Data Figure 5. Guinier approximations of SAXS data obtained for *wt* EnB1 at different concentrations: **a**) 0.69 mg/ml, **b**) 0.34 mg/ml and **c**) 0.14 mg/ml, and EnB1_$\Delta SH_3$ at 2.5 mg/ml (**d**).

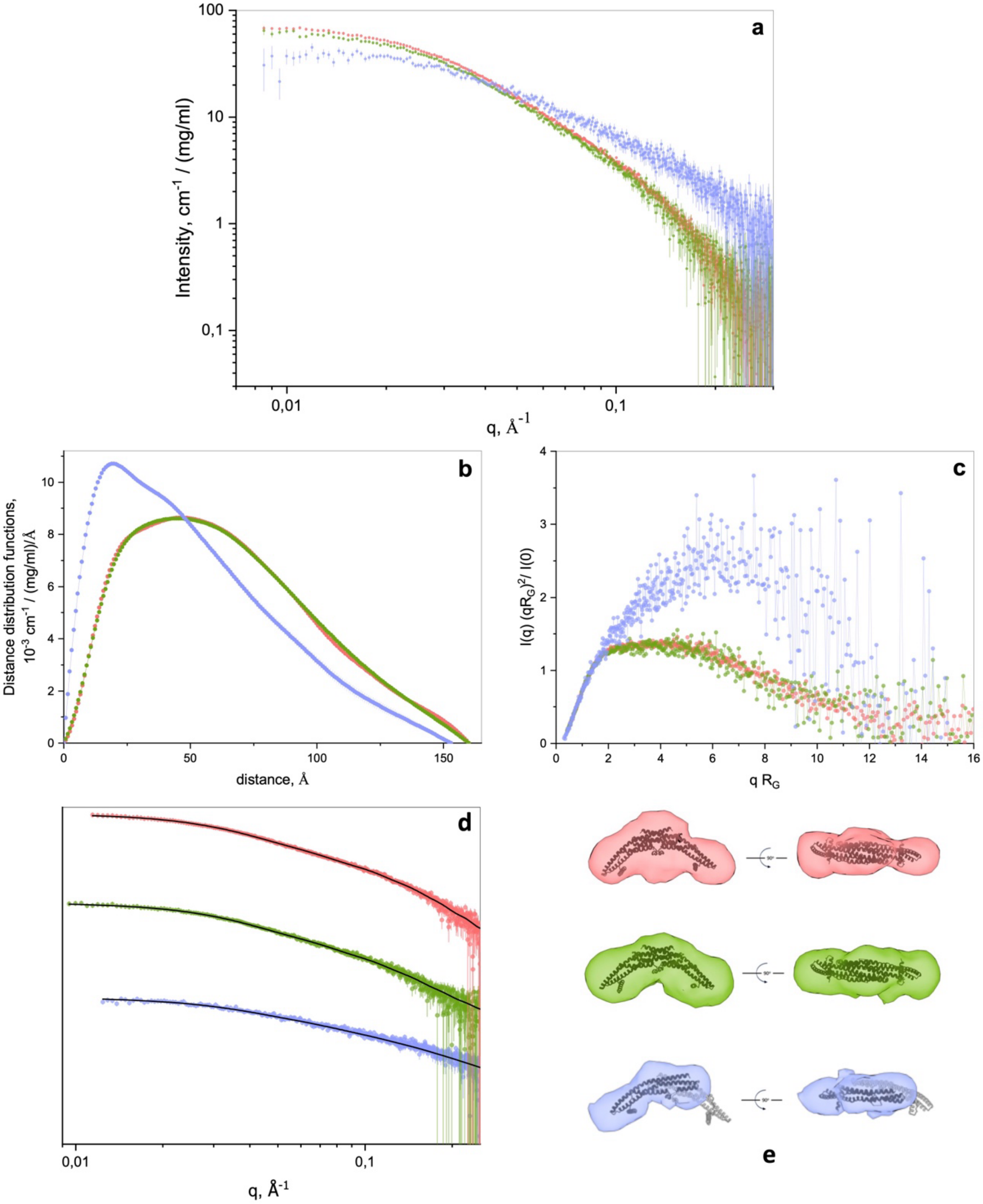

Extended Data Figure 6. SAXS curves for *wt* EnB1, corresponding distance distribution functions and Normalized Kratky plots. **a**) SAXS experimental data of *wt* EnB1 at concentrations 0.69 mg/ml (red), 0.34 mg/ml (green) and 0.14 mg/ml (blue). **b**) Distance distribution functions calculated from SAXS experimental data shown in (**a**). **c**) Normalized Kratky plots obtained from SAXS: I(0) and $R_G$ values were calculated from *P(r).* **d**) Model fits (black lines) of data shown in (**a**) obtained from GNOM calculations and DENSSweb software. **e**) SAXS *ab-initio* reconstructions determined from SAXS data of *wt* EnB1 at different concentrations and comparisons of each with the cryo-EM center dimer atomic structure (black ribbons).

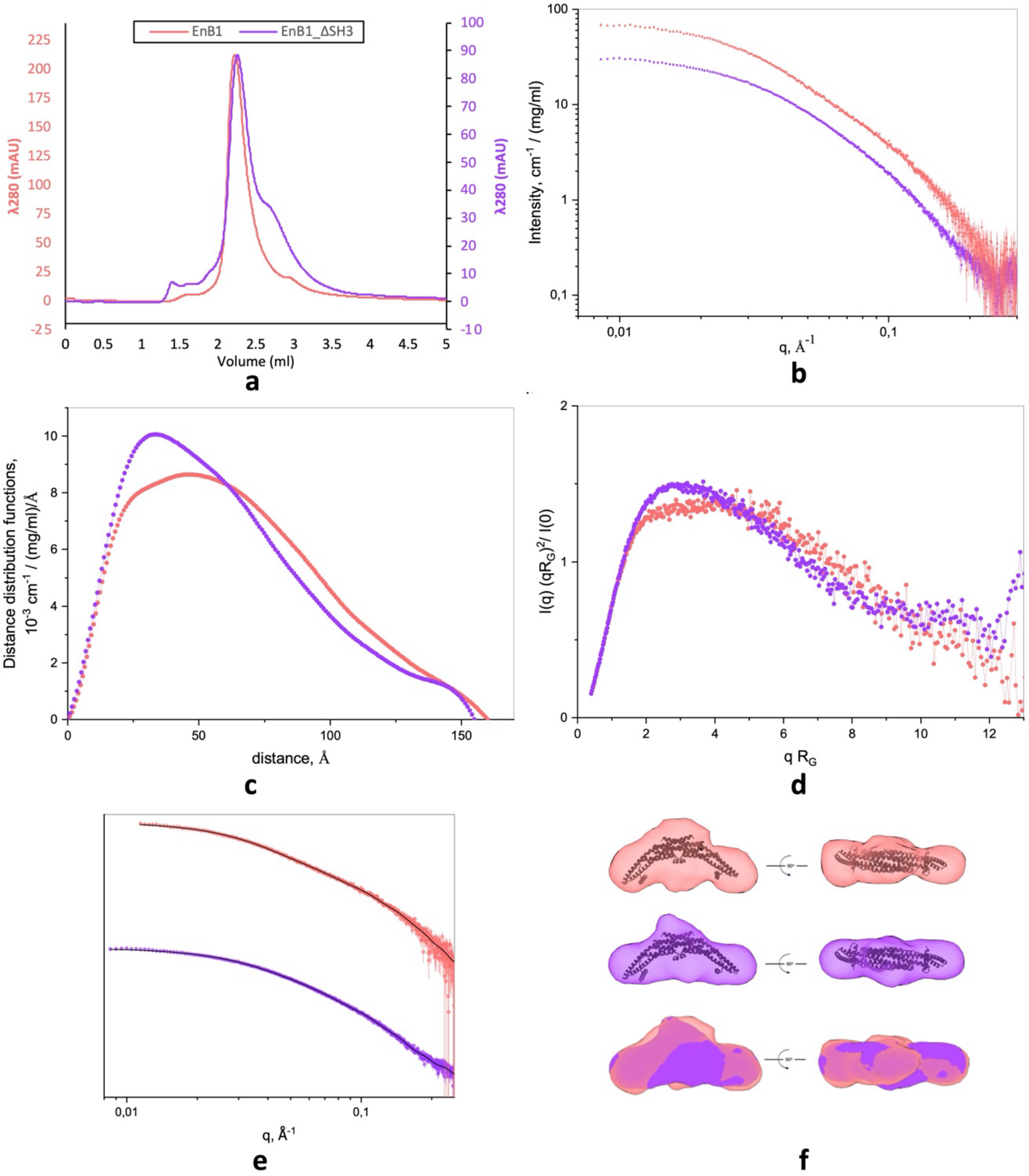

Extended data figure 7. SEC chromatogram and SAXS curves from analysis of EnB1 and EnB1_ΔSH3. **a**) EnB1 (red) and EnB1_ΔSH3 (purple) elute at 2.24 ml and 2.27 ml, respectively, from an AdvanceBio SEC 300Å, 4.6 x 300 mm, 2.7 µm, HPLC column (Agilent). **b**) SAXS experimental data of EnB1 (red) and EnB1_ΔSH3 (purple). **c**) Distance distribution functions calculated from SAXS experimental data shown in (a). **d**) Normalized Kratky plots obtained from SAXS: I(0) and $R_G$ values were calculated from *P(r)*. **e**) Model fits (black lines) for EnB1 and EnB1_ΔSH3 obtained from GNOM calculations and DENSSweb software. Electron density maps obtained for EnB1 and EnB1_ΔSH3, comparison of them with atomic structure (black ribbons). **f**) Comparison of SAXS *ab-initio* reconstructions.

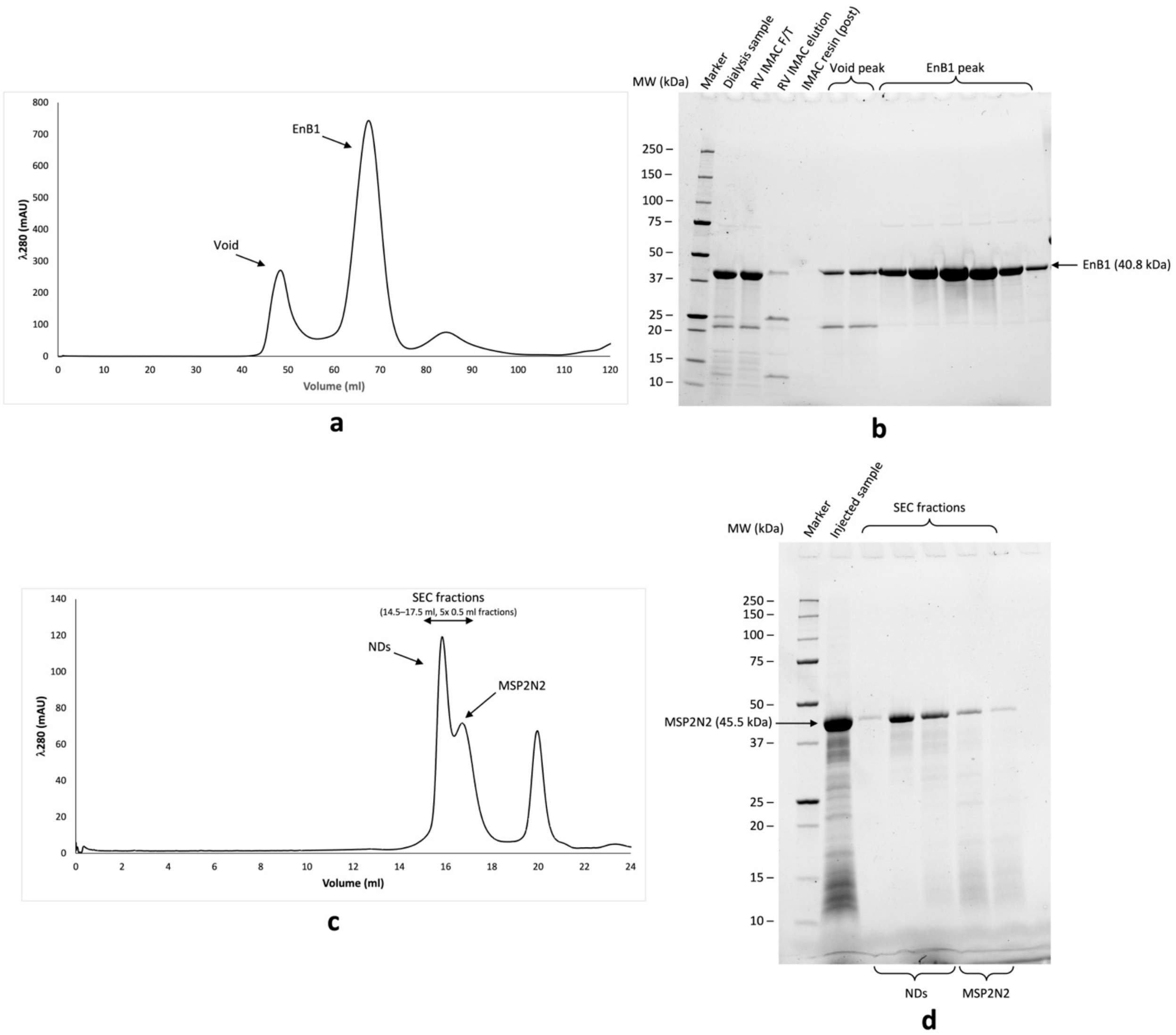

Extended data figure 8. Purification of EnB1 and MSP2N2 NDs. **a**) Representative SEC chromatogram from analysis of EnB1 using a Superdex 200 Increase 16/600 column (GE Healthcare). **b**) Stain-free visualization (Bio-Rad) of SDS-PAGE analysis of SEC run shown in (a). **c**) Representative SEC chromatogram from analysis of MSP2N2 NDs using a Superose 6 10/300 GL (Cytiva) column. **d**) Stain-free visualization of SDS-PAGE analysis of SEC run shown in (**c**).

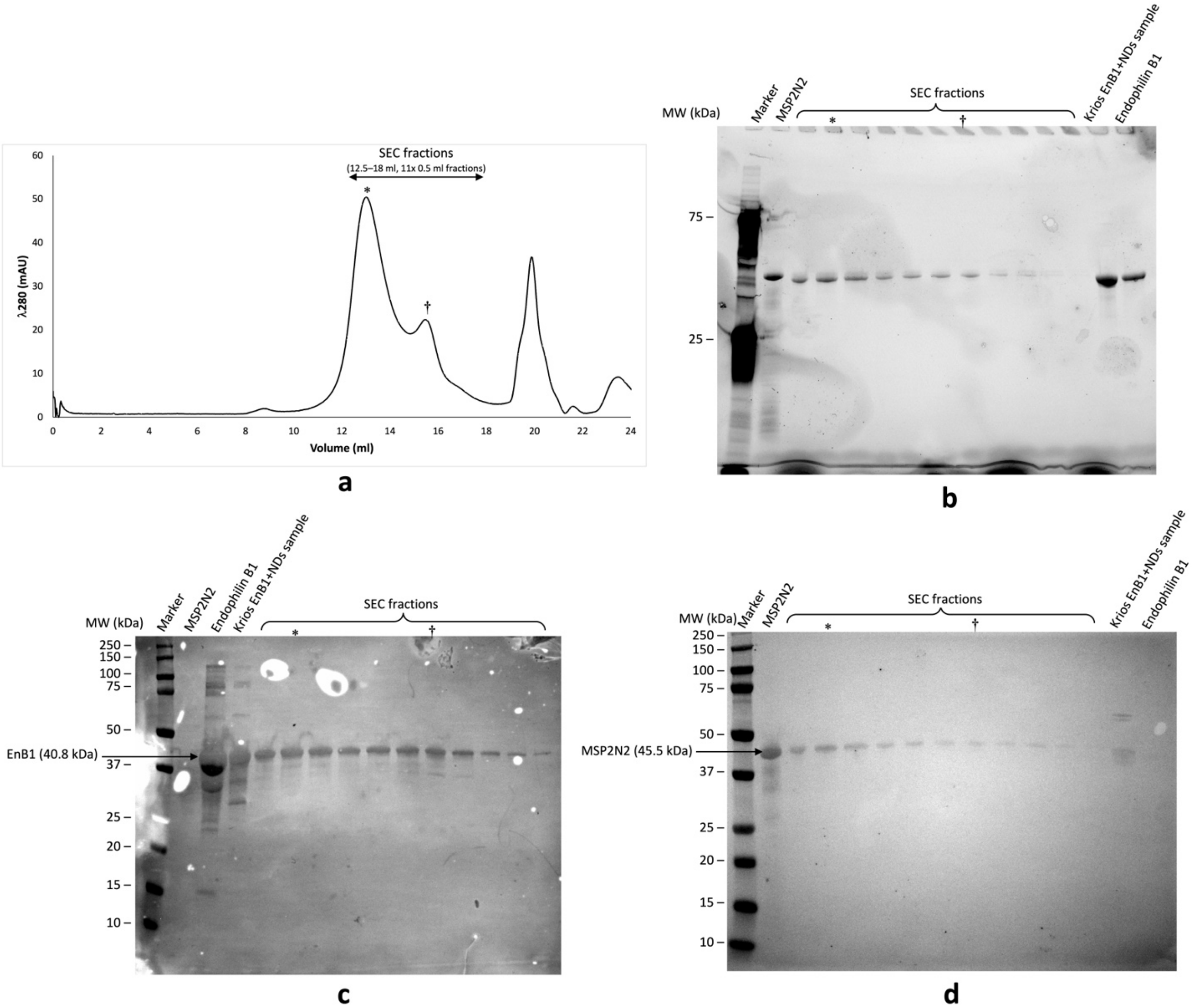

Extended data figure 9. Results from a purification of EnB1 decorated NDs (1:10 MSP2N2: EnB1). **a**) Representative SEC chromatogram from analysis of EnB1 incubated with MSP2N2 NDs using a Superose 6 10/300 GL column. **b**) Stain-free visualization of SDS-PAGE analysis of peak fractions from SEC run shown in (a), purified EnB1 and MSP2N2 and a previous EnB1-decorated NDs sample used for cryo-EM data collection (Krios EnB1+NDs sample). Western blot analyses samples shown in (**b**) using anti-EnB1 (**c**) and anti-polyHis (**d**) primary antibodies and alkaline phosphatase conjugated secondary antibodies. Fractions from the dominant peak are marked with an asterisk (*) contain both EnB1 and MSP2N2. The sample used for the high-resolution Krios data collection also contains both EnB1 and MSP2N2.

### Extended Data Tables

Extended Data Table 1. Cryo-EM data collection, refinement and validation statistics.

| | #1 Endophilin B1 bound to MSP2N2 nanodiscs (consensus map) (EMDB-50981) (PDB 9G2R) | #2 Endophilin B1 bound to MSP2N2 nanodiscs (center dimer focused map) (EMDB-50984) (PDB 9G2U) | #2 Endophilin B1 bound to MSP2N2 nanodiscs (side dimer focused map) (EMDB-50986) (PDB 9G2W) |
|---|---|---|---|
| **Data collection and processing** | | | |
| Magnification | 130,000 | 130,000 | 130,000 |
| Voltage (kV) | 300 | 300 | 300 |
| Electron exposure (e–/Å$^2$) | 40 | 40 | 40 |
| Defocus range (μm) | -1.2 to -2.2 | -1.2 to -2.2 | -1.2 to -2.2 |
| Pixel size (Å) | 0.664 | 0.664 | 0.664 |
| Symmetry imposed | C1 | C1 | C1 |
| Initial particle images (no.) | 5,026,978 | 5,026,978 | 5,026,978 |
| Final particle images (no.) | 273,120 | 273,120 | 273,120 |
| Map resolution (Å) | 3.88 | 3.45 | 3.60 |
| FSC threshold | 0.143 | 0.143 | 0.143 |
| Map resolution range (Å) | 2.98 to 30.00 | 2.98 to 30.00 | 2.98 to 30.00 |
| **Refinement** | | | |
| Initial model used (AlphaFold code) | AF-Q9Y371-F1-model_v4 | AF-Q9Y371-F1-model_v4 | AF-Q9Y371-F1-model_v4 |
| Model resolution (Å) | 3.9 | 3.4 | 3.5 |
| FSC threshold | 0.143 | 0.143 | 0.143 |
| Model resolution range (Å) | 3.26 to 8.96 | 2.98 to 7.38 | 3.00 to 7.02 |
| Map sharpening *B* factor (Å$^2$) | 108.8 | 91.5 | 115.4 |
| Model composition | | | |
| Non-hydrogen atoms | 22620 | 3634 | 3838 |
| Protein residues | 2824 | 456 | 478 |
| Ligands | 0 | 0 | 0 |
| *B* factors (Å$^2$) | | | |
| Protein | 93.50/273.63/156.42 | 93.50/201.89/141.75 | 105.94/273.63/163.37 |
| Ligand | N/A | N/A | N/A |
| R.m.s. deviations | | | |
| Bond lengths (Å) | 0.011 (0) | 0.012 (0) | 0.010 (0) |
| Bond angles (°) | 1.242 (0) | 1.328 (0) | 1.200 (0) |
| Validation | | | |
| MolProbity score | 1.59 | 1.52 | 1.58 |
| Clashscore | 11.85 | 9.87 | 11.68 |
| Poor rotamers (%) | 0.58 | 0.26 | 0.73 |
| Ramachandran plot | | | |
| Favored (%) | 98.28 | 98.66 | 98.09 |
| Allowed (%) | 1.72 | 1.34 | 1.91 |
| Disallowed (%) | 0.00 | 0.00 | 0.00 |

Extended Data Table 2. SAXS data collection and refinement statistics

| | | | | |
|---|---|---|---|---|
| ***(a)* Sample details[1]** | | | | |
| | *Endophilin B1_wt 0.69mg/ml* | *Endophilin B1_wt 0.34mg/ml* | *Endophilin B1_wt 0.14mg/ml* | *Endophilin B1_ΔSH3* |
| Description of sequence | Endophilin-B1 (Q9Y371) from *Homo sapiens* Full-length | | | Truncated (1-306a.a.) |
| Extinction coefficient ε ($M^{-1}$ $cm^{-1}$) | 29130 (280nm) | | | 19160 (280nm) |
| Partial specific volume ($cm^3/g$) | 0,72865 | | | 0.60219 |
| Mean solute and solvent SLD ($10^{-6}$ $Å^{-2}$)[2] | 12.475 / 9.465 | | | 15.094 / 9.465 |
| Mean scattering contrast ($10^{-6}$ $Å^{-2}$)[2] | 3.010 | | | 5.629 |
| Molecular mass (Da)[2] | 40794 | | | 34219 |
| Sample concentration (mg/ml) | 0.69 | 0.34 | 0.14 | 2.5 |
| Solvent composition | 150mM NaCl, 20mM HEPES, 1mM TCEP, 0.5mM DTT, pH 8.1 | | | |
| ***(b)* SAS data collection parameters** | | | | |
| Instrument | ESRF BM29 | | | |
| Wavelength (Å) | 0.9918 | | | |
| Beam geometry | Size: 700 × 700 $\mu m^2$; Sample-to-detector distance: 2.827 m | | | |
| Sample configuration | 1.0 mm-diameter quartz capillary | | | |
| q-measurement range ($Å^{-1}$) | 0.0025 – 0.6 | | | |
| Absolute scaling method | Comparison with scattering from pure $H_2O$ | | | |
| Basis for normalization to constant counts | To transmitted intensity by direct beam counter | | | |
| Exposure time | 15 sec | | | |
| Sample temperature (°C) | 4 | | | |
| ***(c)* Software employed for SAS data reduction, analysis and interpretation** | | | | |
| SAS data averaging and subtraction | PRIMUS from ATSAS 3.2.1 | | | |
| Calculation of ε from sequence | ProtParam: https://web.expasy.org/protparam/ | | | |
| Calculation of values from chemical composition | Peptide Property Calculator: http://biotools.nubic.northwestern.edu/proteincalc.html | | | |
| Calculation of values from chemical composition | SLD calculator web: http://www.refcalc.appspot.com/sld | | | |
| Guinier, *P*(*r*) | GNOM from ATSAS | | | |
| Atomic structure map modelling | DENSSWeb (v 1.7.0): https://denss.ccr.buffalo.edu | | | |
| Molecular graphics | ChimeraX-1.6.1 | | | |
| ***(d)* Structural parameters** | | | | |
| ***Guinier analysis*** | *Endophilin B1_wt 0.69mg/ml* | *Endophilin B1_wt 0.34mg/ml* | *Endophilin B1_wt 0.14mg/ml* | *Endophilin B1_ΔSH3* |
| I(0) ($cm^{-1}$) | 73.13 ± 0.28 | 66.06 ± 0.57 | 43.00 ± 0.75 | 31.25 ± 0.07 |
| $R_G$ (Å) | 50.59 ± 2.34 | 49.67 ± 6.09 | 36.88 ± 9.56 | 46.18 ± 2.62 |
| q $R_G$ – range | 0.5748-1.4567 | 0.4702-1.3158 | 0.4571-1.3367 | 0.4632-1.3461 |
| ***P(r) analysis*** | | | | |
| I(0) ($cm^{-1}$) | 72.26 ± 0.23 | 65.95 ± 0.36 | 45.16 ± 0.65 | 31.42 ± 0.06 |
| $R_G$ (Å) | 51.05 ± 1.70 | 51.04 ± 3.60 | 42.70 ± 6.5 | 47.60 ± 1.1 |
| $d_{max}$ (Å) | 160 | 160 | 153 | 152 |
| q-range ($Å^{-1}$) | 0.0114-0.3009 | 0.0095-0.3009 | 0.0124-0.3009 | 0.0085-0.2390 |
| Total quality estimate (GNOM) | 0.8429 | 0.845 | 0.6489 | 0.552 |
| ***(e)* Atomistic modelling** | | | | |
| | *Endophilin B1_wt 0.69mg/ml* | *Endophilin B1_wt 0.34mg/ml* | *Endophilin B1_wt 0.14mg/ml* | *Endophilin B1_ΔSH3* |
| Method | DENSS 1.7.0 | | | |
| q-range for fitting | 0.0114-0.3009 | 0.0095-0.3009 | 0.0124-0.3009 | 0.0085-0.2390 |
| Correlation score | 0.933 | 0.934 | 0.947 | 0.969 |
| ***(f)* Data and model deposition IDs** | | | | |
| | *Endophilin B1_wt 0.69mg/ml* | *Endophilin B1_wt 0.34mg/ml* | *Endophilin B1_wt 0.14mg/ml* | *Endophilin B1_ΔSH3* |
| | SASDVR4 | SASDVS4 | SASDVT4 | SASDVU4 |
